# Chromatin profiling reveals genome stability heterogeneity in clinical isolates of the human pathogen *Aspergillus fumigatus*

**DOI:** 10.1101/2021.04.19.440431

**Authors:** Ana Cristina Colabardini, Fang Wang, Zhengqiang Miao, Lakhansing Pardeshi, Clara Valero, Patrícia Alves de Castro, Daniel Yuri Akiyama, Kaeling Tan, Luisa Czamanski Nora, Rafael Silva-Rocha, Marina Marcet-Houben, Toni Gabaldón, Taicia Fill, Koon Ho Wong, Gustavo H. Goldman

**Affiliations:** Faculdade de Ciências Farmacêuticas de Ribeirão Preto, Universidade de São Paulo, Ribeirão Preto, São Paulo, Brazil; Faculty of Health Sciences, University of Macau, Macau SAR of China; Genomics, Bioinformatics and Single Cell Analysis Core, Faculty of Health Sciences, University of Macau, Macau SAR of China; Institute of Translational Medicine, Faculty of Health Sciences, University of Macau, Macau SAR of China; Instituto de Química, Universidade Estadual de Campinas, Campinas, São Paulo, Brazil; Faculdade de Medicina de Ribeirão Preto, Universidade de São Paulo, Ribeirão Preto, São Paulo, Brazil; Barcelona Supercomputing Centre (BSC-CNS). Jordi Girona, Barcelona, Spain; Institute for Research in Biomedicine (IRB Barcelona), The Barcelona Institute of Science and Technology, Baldiri Reixac, Barcelona, Spain; Catalan Institution for Research and Advanced Studies (ICREA), Barcelona, Spain

**Keywords:** *Aspergillus fumigatus*, chromatin profiling, histone modification, transposon, secondary metabolites, genome stability

## Abstract

Invasive Pulmonary aspergillosis is a life-threatening infection in immunosuppressed patients caused by the filamentous fungus *Aspergillus fumigatus*. Chromatin structure regulation is important for genome stability maintenance and has the potential to lead to genome rearrangements driving differences in virulence and pathogenesis of different *A. fumigatus* isolates. Here, we compared the chromatin activities of the most investigated clinical isolates Af293 and CEA17 and uncovered striking differences in the number, locations and expression of transposable elements. We found evidence for higher genome instability in Af293 as compared to CEA17 and identified a spontaneous Af293 variant that exhibits gross chromosomal alterations including the loss of a 320 kb long segment in chromosome VIII and the amplification of a biosynthetic gene cluster. As a consequence of these re-arrangements, the variant shows increased secondary metabolites production, growth and virulence. Our work emphasizes genome stability heterogeneity as an evolutionary driver of *A. fumigatus* fitness and virulence.

## Introduction

Invasive Pulmonary Aspergillosis (IPA) is a life-threatening infection in immunosuppressed patients (van de Veerdonk *et al*., 2017; Latgé and Chamilos, 2019) that is caused mainly by the filamentous fungus *Aspergillus fumigatus*, a ubiquitous soil inhabitant that can utilize a wide variety of organic substrates and adapt well to a broad range of environmental conditions (Sugui *et al*., 2015). *A. fumigatus* produces asexual conidia that readily become airborne and can survive a broad range of temperature and water availability (Croft *et al*., 2016). Although the environmental populations of *A. fumigatus* are genotypically diverse (Alshareef and Robson, 2014), several *A. fumigatus* strains are able to infect immunosuppressed patients (Latgé and Chamilos, 2019). Once inhaled by a human host, *A. fumigatus* conidia penetrate deep in the alveoli, where they must survive the surveillance of the host immune system (Dagenais and Keller, 2009; van de Veerdonk *et al*., 2017; Latgé and Chamilos, 2019). Alveolar macrophages kill conidia and germlings within the phagolysosome by producing reactive oxygen species (ROS) and phagolysosomal acidification (Margalit *et al*., 2015), while neutrophils can attach to germlings and release granules containing a variety of antimicrobial compounds (Heinekamp *et al*., 2015; Margalit *et al*., 2015). If the invader succeeds in establishing an infection in a host who is then subjected to an antifungal treatment, *A. fumigatus* hyphae must also survive the fungicidal and/or fungistatic effects of the antifungal drug(s) in order to persist inside the host (Elefanti *et al*., 2013). Thus, it is essential that *A. fumigatus* can adapt its physiology to the changing conditions during the course of infection to thrive as a pathogen (Buscaino, 2019).

Genomic plasticity is a key factor for the success of pathogenic fungi, as it can reprogram transcriptional events and/or generate variability that allow organisms to adapt to different environments and to improve their arsenal of defences against the host (Bignell *et al*., 2016; Buscaino, 2019). Fungal variants with genotypic diversity were generated under mitotic (Seidl and Thomma, 2014; Gusa and Jinks-Robertson, 2019) or meiotic (O’Gorman *et al*., 2009) processes during development in nature, resulting in isolates with genetic differences ranging from single-nucleotide polymorphisms to chromosome rearrangements (Ballard *et al*., 2018; Dos Reis *et al*., 2018). It is known that chromosome rearrangements are driven by long repeat sequences in the fungal pathogen *Candida albicans* (Todd *et al*., 2019; Dunn and Anderson, 2019) and subjected to epigenetic control (Freire-Benéitez *et al*., 2016a), however, the regulatory mechanism of this process is still not fully understood. Epigenetic regulation is coordinated by chromatin, which consists of histone, non-histone proteins and RNA that package the DNA molecules (McGinty and Tan, 2015). Histone tails are prone to various post-translational modifications (PTM) that control the chromatin status and activity (Hammond *et al*., 2017). Euchromatin, which is generally composed of gene-rich, non-repetitive DNAs, and heterochromatin, with low gene density and enriched with repetitive DNA, are usually marked by different modifications on histone H3, such as trimethylation of lysines 4 and 9, respectively (Howe *et al*., 2017; Allshire and Madhani, 2018). Among these, the centromeric and telomeric DNA repeats have important roles in kinetochore function (Uhlmann, 2016) and protection of chromosome ends (O’Sullivan and Karlseder, 2010), respectively, while subtelomeres contain repetitive sequences and silent genes intermingled with conditional active genes, such as those for secondary metabolites (SM) production (Ellahi *et al*., 2015; Rutledge and Challis, 2015; Pfannenstiel and Keller, 2019). Several histone PTMs were described in filamentous fungal species, such as *A. nidulans* (Gacek-Matthews *et al*., 2016; Nützmann *et al*., 2013), *A. oryzae* (Maeda *et al*., 2017; Pham *et al*., 2015), *A. flavus* (Lan *et al*., 2016), *Neurospora crassa* (Adhvaryu *et al*, 2005), *Fusarium fujikuroi* (Janevska *et al*., 2018; Niehaus *et al*., 2016) and *F. graminearum* (Kong *et al*., 2018) with distinct types and functions. However, the *A. fumigatus* histone PTMs regulation and their role in virulence and pathogenicity are not known.

Transposable elements (TEs) play a particular role in genome evolution through the generation of variability and genetic instability (Casacuberta and Gonzalez, 2013). TEs are repeated sequences that can multiply and/or move to distant genomic locations, generating a myriad of effects that include gene duplication or inactivation, disruption of regulatory regions, generation of alternative splicing variants and spreading of epigenetic silencing. Furthermore, TEs can passively act as substrate for ectopic recombination and aberrant transposition events (Casacuberta and Gonzalez, 2013). Two classes of TEs are identified in cells: i) Class I TEs are retrotransposons that use an intermediary RNA molecule and a reverse transcriptase to transpose, while ii) class II TEs are DNA transposons that encode transposases to directly mobilize DNA repeats (Kapitonov and Jurka, 2008). TEs are normally silenced by heterochromatin (Allshire and Madhani, 2018) and can be activated under stress or by the disruption of these supressing mechanisms (Casacuberta and Gonzalez, 2013). It has been shown that the acquired SM gene clusters in *Aspergillus* species are flanked by TEs, implying that TEs activity contributed to the acquisition of these important virulence factors (Lind *et al*., 2017) and potentially other genomic changes. Besides TEs, genetic and genomic changes can also originate from cell segmentation for strain adaptation (Buscaino, 2019), which is best examined in *C. albicans* (Selmecki *et al*., 2006; Forche *et al*., 2009; Forche *et al*., 2011; Rustchenko *et al*., 1994; Janbon *et al*., 1998). *A. fumigatus* genome contains abundance of TEs (Fedorova *et al*., 2008; Bignell *et al*., 2016) and high heterogeneity across different *A. fumigatus* isolates in drug susceptibility, nutrients acquisition and virulence has been reported (Alshareef and Robson, 2014; Abdolrasouli *et al*., 2015; Kowalski *et al*., 2016; Knox *et al*., 2016; Ries *et al*., 2019; Dos Santos *et al*., 2020). Hence, it is interesting to understand whether and how those elements affect the physiology and adaptation of this pathogen to diverse environmental pressures during its evolution.

Af293 and CEA17 (a CEA10 derivative) are the two most common *A. fumigatus* clinical isolates used in the laboratory as references (Nierman *et al*., 2005; Fedorova *et al*., 2008). However, they bear significant differences in their physiology and virulence (Keller, 2017). Recently, many factors, such as carbon and nitrogen metabolism and response to low oxygen stress in the lung, have been shown to be heterogeneous across those and other clinical isolates, which consequently have an impact in the host colonization ability (Abdolrasouli *et al*., 2015; Knox *et al*., 2016; Kowalski *et al*., 2016; Lind *et al*., 2017; Ries *et al*., 2019; Dos Santos *et al*., 2020). Genomic comparisons have shown that Af293 and CEA10 have 98% of overall genomic identity with the most variable regions located within 300 kb from the chromosomal telomeres (Fedorova *et al*., 2008). Af293 and CEA10 unique regions include around 200 genes each, and the majority of unique genes are clustered together in blocks ranging from 10 to 400 kb (Rokas *et al*., 2007, Fedorova *et al*., 2008). However, the effects of chromatin structure and function are still largely unexplored in *A. fumigatus*. To address whether epigenetic factors also contribute to the variability of *A. fumigatus* isolates, we compared the well-established active (H3K4me3) and repressive (H3K9me3) histone codes in Af293 and CEA17. This work shows that the modification pattern of the histone marker H3K4me3 that is associated with active gene expression is similar between the two isolates, while the genomic regions marked by the repressive heterochromatic marker H3K9me3 are variable. Furthermore, we detected key differences in TEs distribution between the two isolates, with Af293 having a significantly higher number of TEs from the LINE family. Consistently, genome instability of Af293, but not of CEA17, was detected with chromosomal changes observed in isolates from different laboratories, including ours. We isolated an Af293 variant that had a spontaneous deletion of 320 kb at the right arm of the chromosome VIII and an amplification of 20 genes which are originally located at the left arm of the same chromosome, and both deletion and amplification sites were adjacent to LINE TEs. Genomic, transcriptomic and phenotypic characterization of the Af293 variant revealed unexpected effects on its growth fitness, mycotoxin production and virulence. Overall, our results have provided the first chromatin evidence that genome stability heterogeneity could play a role in *A. fumigatus* fitness within the host and hence, this work has profound importance in the comprehension of the fungal pathogen environmental adaptation.

## Results

### Active *A. fumigatus* genes undergo nucleosome depletion at promoters and H3K4me3 modification at 5’ coding regions

In order to compare chromatin activities of the two *A. fumigatus* reference strains Af293 and CEA17, we performed Chromatin Immunoprecipitation coupled to high throughput sequencing (ChIP-seq) to map genome wide nucleosome occupancy (histone H3) and histone H3 K4 trimethylation (H3K4me3) that associates with active transcription. Input and immunoprecipitated DNA samples were sequenced using the Illumina HiSeq2500 platform and the sequencing reads were aligned to the respective Af293 or CEA17 genome. It is noteworthy that the CEA17 reference genome is only partially assembled composing of 55 non-overlapping contigs (Fedorova *et al*., 2008), as compared to eight full chromosomal assemblies in the Af293 reference genome (Nierman *et al*., 2005; Ronning *et al*., 2005). The Af293 reference also has relatively much better gene model and function annotations. Therefore, we explored the feasibility of using the Af293 reference genome for CEA17 data. Considering the fact that there are some minor differences between the two genomes (Fedorova *et al*, 2008), we mapped the CEA17 data to both CEA17 and Af293 reference genomes for a comparison and found only a marginal difference in terms of the mapping percentage (97.9% and 95.1% of CEA17 reads mapping to the CEA17 and Af293 reference genome, respectively) (Supplementary Table 1). This suggests that for most parts the use of Af293 reference genome for CEA17 data would have a small effect, if any, on the results and data interpretation. In light of this, results based on mapping to the Af293 genome will be presented, except for analyses whereby genome structure matters (in these cases the CEA17 reference is used and duly indicated).

Consistent with observations in other eukaryotes, promoters (between −500 to +100 bp with respect to annotated transcription start sites [TSS]) of *A. fumigatus* protein coding genes were found depleted for nucleosomes (Figure 1a), while H3K4me3 modifications were enriched at the first half of the open reading frame (ORF) of many genes (Figure 1b). To understand the relationship between promoter nucleosome occupancy, H3K4me3 modification and transcriptional activity, we also performed RNA sequencing (RNA-seq) for the two isolates under the same growth conditions, and correlated nucleosome and H3K4me3 levels with mRNA abundance. We divided Af293 annotated genes into 5 groups based on their gene expression levels determined by RNA-seq from the most (1^st^ 2000) to the least expressed (5^th^ 1825) (Figure 1c, top panel). ChIP-seq analysis of histone H3 occupancy revealed that the two most expressed gene groups had the lowest H3 accumulation at the TSSs in Af293 (Figure 1c, middle panel), which suggests that nucleosomes were depleted at these promoters (*i.e.*, a permissive state for transcription). On the other hand, the expression levels of genes in the 3^rd^, 4^th^ and 5^th^ groups were inversely proportional to their promoter H3 levels (Figure 1c, middle panel), reflecting a general negative relationship between promoter nucleosome density and transcriptional activity, consistent with the well-established role of nucleosome occupancy and positioning in controlling transcriptional activation.

**Figure 1.**
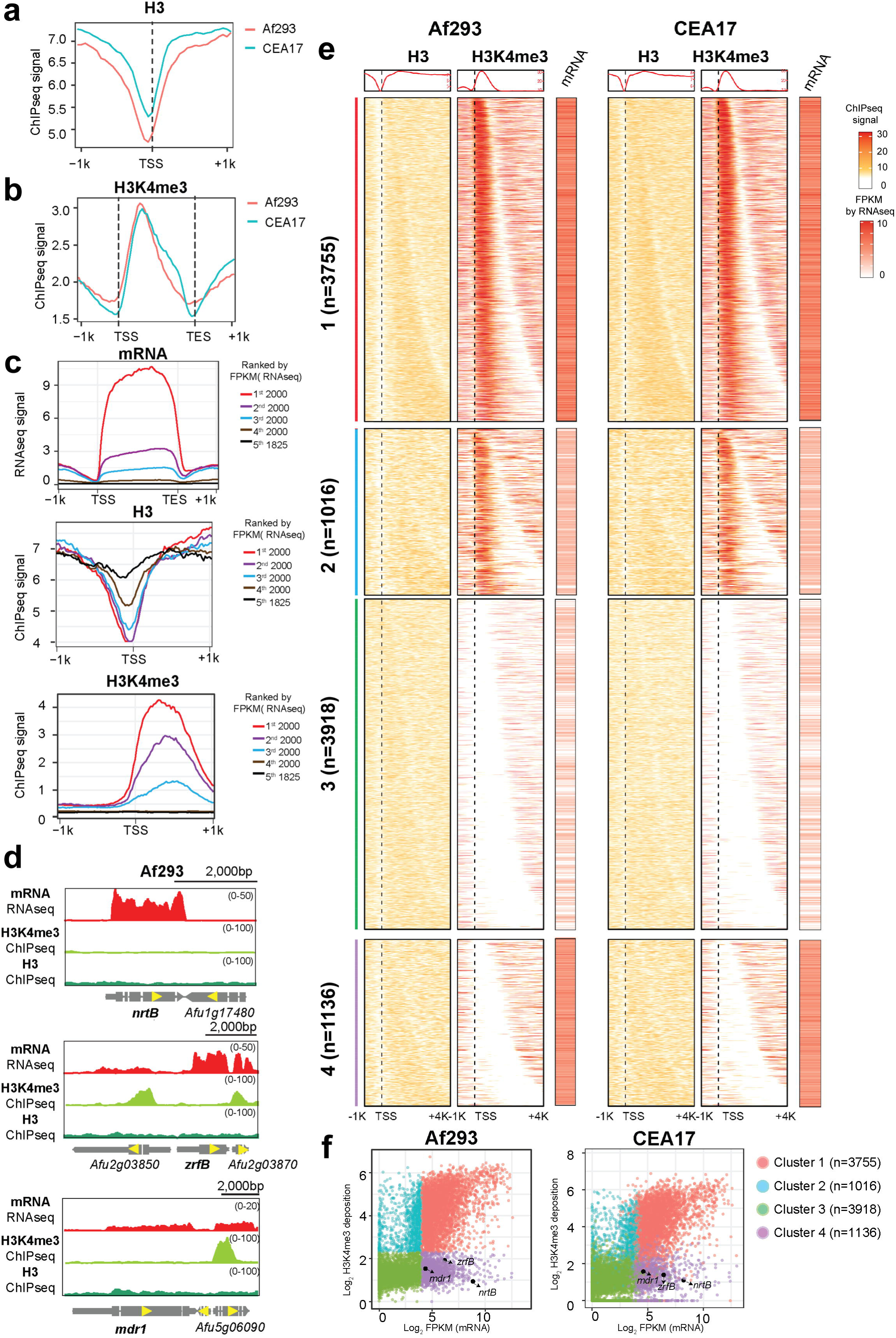
The nucleosome depletion at promoters and H3K4me3 modification at 5’ coding regions of two *A. fumigatus* isolates Af293 and CEA17. a, Line plots showing the H3 deposition within 1kb of TSS region of genes genome wide. b, Line plots showing the H3K4me3 deposition within 1kb of CDS region of genes genome wide. c, Line plots showing the mRNA level (top), and deposition of H3 (middle) and H3K4me3 (bottom) within 1kb of CDS or TSS regions of genes genome wide. Gene order is ranked by mRNA level from high to low. d, Genome browser screenshots showing mRNA and H3K4me3 level at selected genes. H3 level was used as control. e, Heatmap plots showing the mRNA accumulation and deposition of H3 and H3K4me3 in Af293 and CEA17 genes genome wide classified as four clusters. f, Scatter plots showing the mRNA and H3K4me3 levels of the four cluster genes as shown in (e).

In contrast, H3K4me3 modification is positively correlated with transcriptional activity in which the most expressed genes had high levels of H3K4me3 modification at the 5’ end of ORFs, and the levels reduced proportionally in the second and third groups (Figure 1c, bottom panel). The correlated gene expression and H3K4me3 suggests a positive association between H3K4me3 modification and transcriptional activity in Af293, agreeing with published literature of other eukaryotes (Howe *et al*., 2017). For the CEA17 isolate, the same correlations of H3 and H3K4me3 with transcriptional activity (mRNA levels by RNA-seq) were observed, regardless of which reference genome was used for the analysis (Supplementary Figure 1a). Global gene expression levels (*e.g.*, FPKM from RNA-seq) and H3K4me3 ChIP-seq profiles were also highly correlated between the Af293 and CEA17 isolates (Supplementary Figure 1b). Taken together, these observations indicate negative and positive effects of promoter nucleosome and H3K4me3 modification on transcriptional activity, respectively, in *A. fumigatus* and these chromatin regulations are general in the two different clinical isolates.

### Many expressed genes are devoid of H3K4me3 modification

In the course of data inspection on a genome-browser, we noted many well-expressed genes (*e.g. nrtB*, *zrfB*, and *mdr1*) without H3K4me3 modification at their coding regions for both Af293 (Figure 1d) and CEA17 (Supplementary Figure 1c), suggesting a potentially interesting difference in the mechanistic relationship between transcription and H3K4me3 modification. To systematically analyse this, four gene clusters were classified according to the level of H3K4me3 modification and mRNA expression levels (Figure 1e). Genes in cluster 1 (n = 3755) had high H3K4me3 modification and gene expression levels (Figure 1e, f). On the other hand, genes in cluster 2 (n = 1016) had relatively lower mRNA levels when compared to cluster 1, but significant amount of H3K4me3 ChIPseq signals could still be observed (Figure 1e, f). On the other hand, cluster 3 genes (n = 3918) had low or background levels of H3K4me3 and mRNA, which presumably is indicative of limited transcriptional activities (Figure 1e, f). Interestingly, a large group of well-expressed genes (n = 1136; cluster 4) showed a lack of H3K4me3 modification, similar to what was observed at the genome-browser (Figure 1e, f). The same observation was found for CEA17 (Figure 1e, f), suggesting that this is a general phenomenon for *A. fumigatus*.

Gene Ontology (GO) enrichment analysis revealed that cluster 1 genes are related to primary metabolic processes such as translation, cellular component organization and biosynthesis (Supplementary Figure 1d, Supplementary Data 1), while genes in cluster 2 were mainly associated to DNA metabolism, DNA repair, tRNA processing, autophagy and melanin biosynthesis (Supplementary Figure 1d, Supplementary Data 1). As expected, the lowly or non-expressed genes in cluster 3 were highly enriched with the secondary metabolism processes (Supplementary Figure 1d, Supplementary Data 1), although cell wall remodelling genes were also present in this cluster (Supplementary Data 1). Interestingly, those genes with an incongruous H3K4me3 modification and gene expression levels (cluster 4) encode proteins involved in transmembrane transport functions such as carbohydrate, ion, ammonium and drug transporters (Supplementary Figure 1d, Supplementary Data 1), suggesting that genes involved in transport processes may have an atypical mechanistic relationship between transcription and H3K4me3 modification.

### Af293 and CEA17 H3K9me3 modification profiles are similar at (sub)telomeric regions but differ at many chromosomal locations

After evaluation of the active chromatin marker, we also compared the histone modification H3K9me3 using ChIP-seq to investigate the heterochromatin situation in the two isolates. At the genome browser level, high enrichments of H3K9me3 modification were found at all ends (*i.e.* telomeres) (Figure 2a), similar to what had been reported in other eukaryotes such as *Schizosaccharomyces pombe* (Nakayama *et al*. 2001; Yamada *et al*. 2005) and *Neurospora crassa* (Rountree and Selker, 2010) but unlike *Candida albicans* (Freire-Benéitez *et al*., 2016; Price *et al*., 2019) and *Saccharomyces cerevisiae* (O’Kane and Hyland, 2019). The modification patterns at these chromosomal regions are highly similar between Af293 and CEA17 when analysed using the Af293 reference genomes (Figure 2a, Supplementary Figure 2a) except for the region genome reference matters (*e.g.* the subtelomeric region with low CEA17 reads when alignment to chromosome I of Af293 genome) (Fedorova *et al*., 2008). In order to rule out any other potential mapping artefact due to genome differences, we then mapped the CEA17 data to its own reference genome. While a full analysis on all centromeric and telomeric regions is not possible for CEA17 data due to its partial assembled genome and poor annotation, H3K9me3 peaks were observed at both ends of the largest scaffolds (Scf_1, 2, 3 and 4) (Figure 2b, Supplementary Figure 2b), which presumably are full chromosomal assembly (Fedorova *et al*., 2008) corresponding to the Af293 chromosomes 1, 2, 3 and 5, respectively. Therefore, H3K9me3 modified nucleosomes are deposited at chromosome ends (*i.e.*, subtelomere or telomere in both Af293 and CEA17), which presumably are full chromosomal assembly (Fedorova *et al*., 2008) corresponding to the Af293 chromosomes 1, 2, 3 and 5, respectively. Correspondingly, the expression of those subtelomeric genes have much lower gene expression (mRNA) and H3K4me3 level under the primary culture conditions (Figure 2c), and approximately 75% of those genes belong to beforementioned cluster 3 (Supplementary Figure 2c), reinforcing the role of transcriptionally silent heterochromatin H3K9me3 at telomeres in *A. fumigatus*.

**Figure 2.**
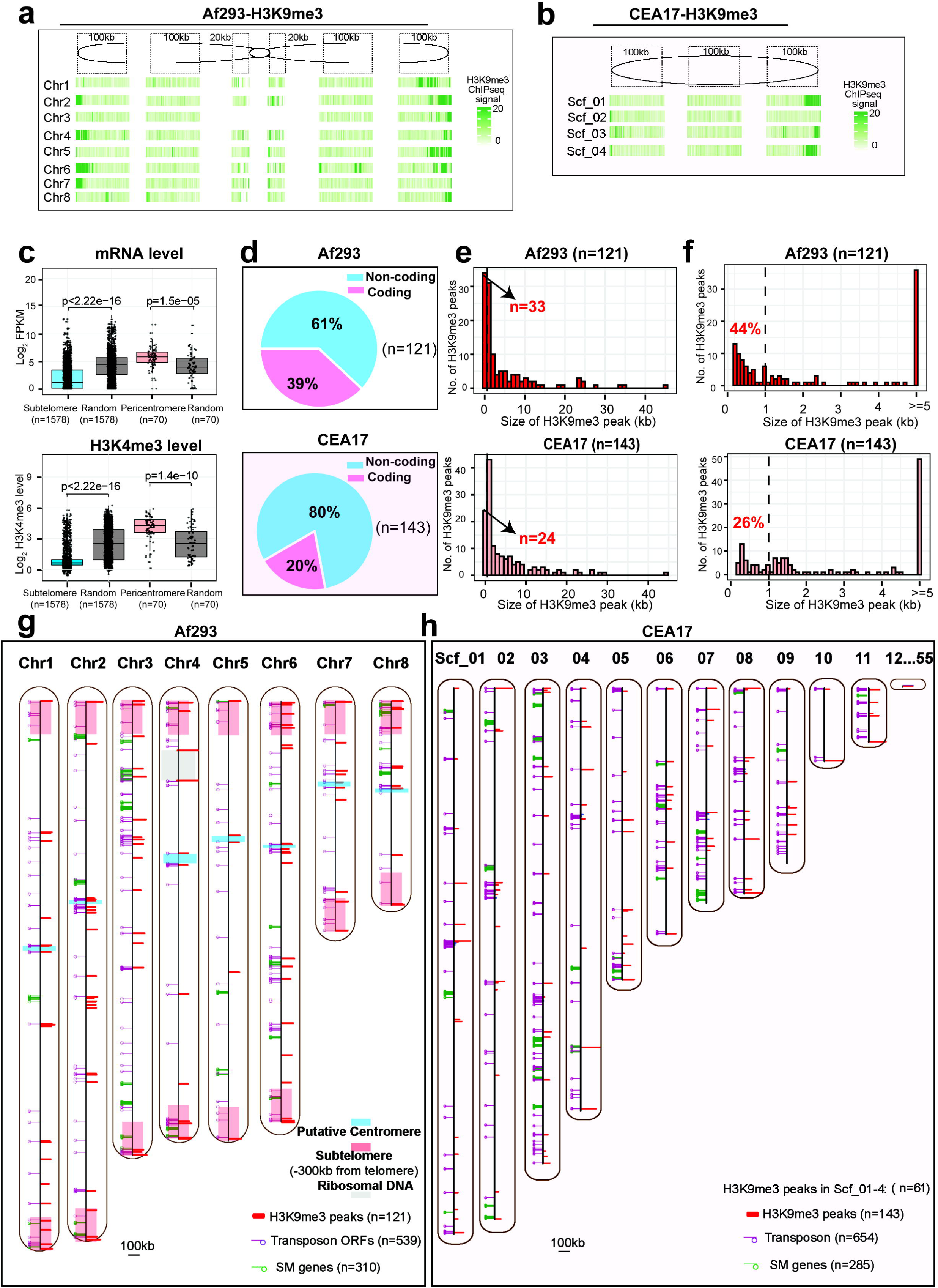
Different H3K9me3 modification profiles in Af293 and CEA17. a, A heatmap plot showing the H3K9me3 levels in Af293 genome. Each bin represents 200bp; 100kb nearby chromosome end, 20kb pericentromere, and randomly selected 100kb regions were plotted for each chromosome. b, A heatmap plot showing the H3K9me3 levels in CEA17 four big contigs. The pink shade was plotted as shown the mapping was performed to CEA17 genome reference. c, Box plots showing the mRNA (top panel) and H3K4me3 (panel) level of selected genes in Af293. d, A sector diagram showing the genome wide distribution of H3K9me3 peaks called by MACS2 in Af293 and CEA17. e-f, Histogram plots showing the size of H3K9me3 peaks in Af293 and CEA17 isolates. g-h, A scheme representing the genome wide location of centromeres (blue), subtelomeres (pink), rDNAs loci (grey), H3K9me3 binding peaks (red), transposon genes (purple) and SM biosynthetic genes (green) along the 8 chromosomes of Af293 (g) and 11 contigs of CEA17 genome (h).

Further, we investigated the chromatin structure in/near centromere loci in Af293. Putative centromere loci were called according to the genomic DNA gap from sequencing output reads (Supplementary Table 2). Relatively high H3K9me3 signal was also observed at pericentromere loci from genome browser investigation (Figure 2a and Supplementary Figure 2d). Interestingly, high amount of H3K4me3 and gene expression level were also detected for those pericentromere genes (*i.e.,* the nearby 5 genes at both side of centromere) (Figure 2c), among which, 85% of were belong to beforementioned cluster 1 (Supplementary Figure 2e). The result is consistent as *C. albicans* that the pericentromere beared both euchromatin and heterochromatin features. Overall, our results showed H3K9me3 modified nucleosomes represent heterochromatin and have similar deposition at (sub)telomeres in both Af293 and CEA17.

To systematically identify heterochromatin regions (*i.e.* H3K9me3 modified regions) throughout the genome, we applied MACS2 broad peaks calling method on the H3K9me3 ChIP-seq data of the two isolates mapped to their respective reference genome. A total of 121 and 143 H3K9me3 peaks were obtained for Af293 and CEA17, respectively (Supplementary Data 2). Most of them were located in non-coding regions (Figure 2d, Supplementary Data 2). Interestingly, the size of peaks (*i.e.*, the span of H3K9me3 modified region) varied significantly within the isolates, ranging from 0.2 to 45 kb (Figure 2e). Specifically, 88 and 119 peaks in Af293 and CEA17, respectively, spread across broad regions (>500bp), while 33 and 24 peaks had narrower modified regions (<500bp) (Supplementary Data 2). Interestingly, the peak sizes differed significantly between the two isolates with Af293 having relatively shorter H3K9me3 modified regions (*e.g.* <500 bp) than CEA17 (Figure 2f), indicating dissimilar heterochromatin patterns in the two isolates.

We next assigned the H3K9me3 peaks to chromosomes. Of the 121 Af293 H3K9me3 peaks, 20 were found near (*e.g.* within 10 kb) centromeres and telomeres and 21 were located within 300 kb from chromosome ends (Figure 2g, Supplementary Data 2), which are commonly referred as the subtelomeric regions (McDonagh *et al*., 2008). It is noteworthy that the majority of peaks were actually found distributed across chromosome bodies (*i.e.*, not at telomeric and subtelomeric regions). Although the analysis for CEA17 was limited by its incomplete genome assembly, H3K9me3 peaks were clearly observed at telomeric (n = 8), subtelomeric (n = 8) as well as internal regions (n = 45) of the four largest scaffolds (Scf_1, 2, 3 and 4). More importantly, with the exception for a few regions, the distribution of H3K9me3 peaks on the corresponding chromosome bodies of the two isolates was somewhat different (Figure 2g, h and Supplementary Figure 2b). Therefore, these results reveal dissimilarities in the pattern of H3K9me3 deposition on Af293 and CEA17 genomes.

### Most BGCs are not modified by the H3K9me3 heterochromatic modification in *A. fumigatus*

One of the major classes of genes subjected to H3K9me3 regulation (*i.e.*, transcriptional silencing) is the secondary metabolite (SM) biosynthetic genes, which are often arranged in clusters (refer to as BGCs hereafter) (Pfannensiel and Keller, 2019). Therefore, it is possible that the differences in H3K9me3 modifications in the two isolates are related to differences in SM BGCs regulation and/or their genomic locations. However, as shown previously, the genomic locations of most SM BGCs on the four largest scaffolds of CEA17 (n = 17) were similar to the corresponding chromosome (Chr1, 2, 3, 5) of Af293 (n = 19), with the exception of those BGCs (BGCs 9-16) whose genomic arrangement was inverted between CEA17 Scf_3 and Af293 Chr3 (Supplementary Figure 3a) and the BGCs (BGCs 27 and 33) that were non-located at corresponding chromosomes in two isolates (Supplementary Table 3) (Fedorova *et al*., 2008). To better analyse the situation, we mapped all identified H3K9me3 peaks to SM BGCs in the two isolates. There are 31 common SM BGCs in both Af293 and CEA17, and Af293 genome contains two additional SM BGCs (BGC 1 and 4) that are not present in CEA17 (Lind *et al*., 2017) (Supplementary Table 3). Surprisingly, we found only two H3K9me3 peaks located within SM BGCs for both Af293 (BGCs 10 and 16) (Figure 3a) and CEA17 (BGCs 16 and 21) (Supplementary Figure 3b); even though the ChIP-seq experiment was carried out under the conditions in which most of the SM BGCs are silent. Moreover, the peaks found at SM BGCs only spanned a relatively small and gene-free region (*e.g.,* about 1-8% of the size of BGC on average) (Figure 3a and Supplementary Figure 3b, c).

**Figure 3.**
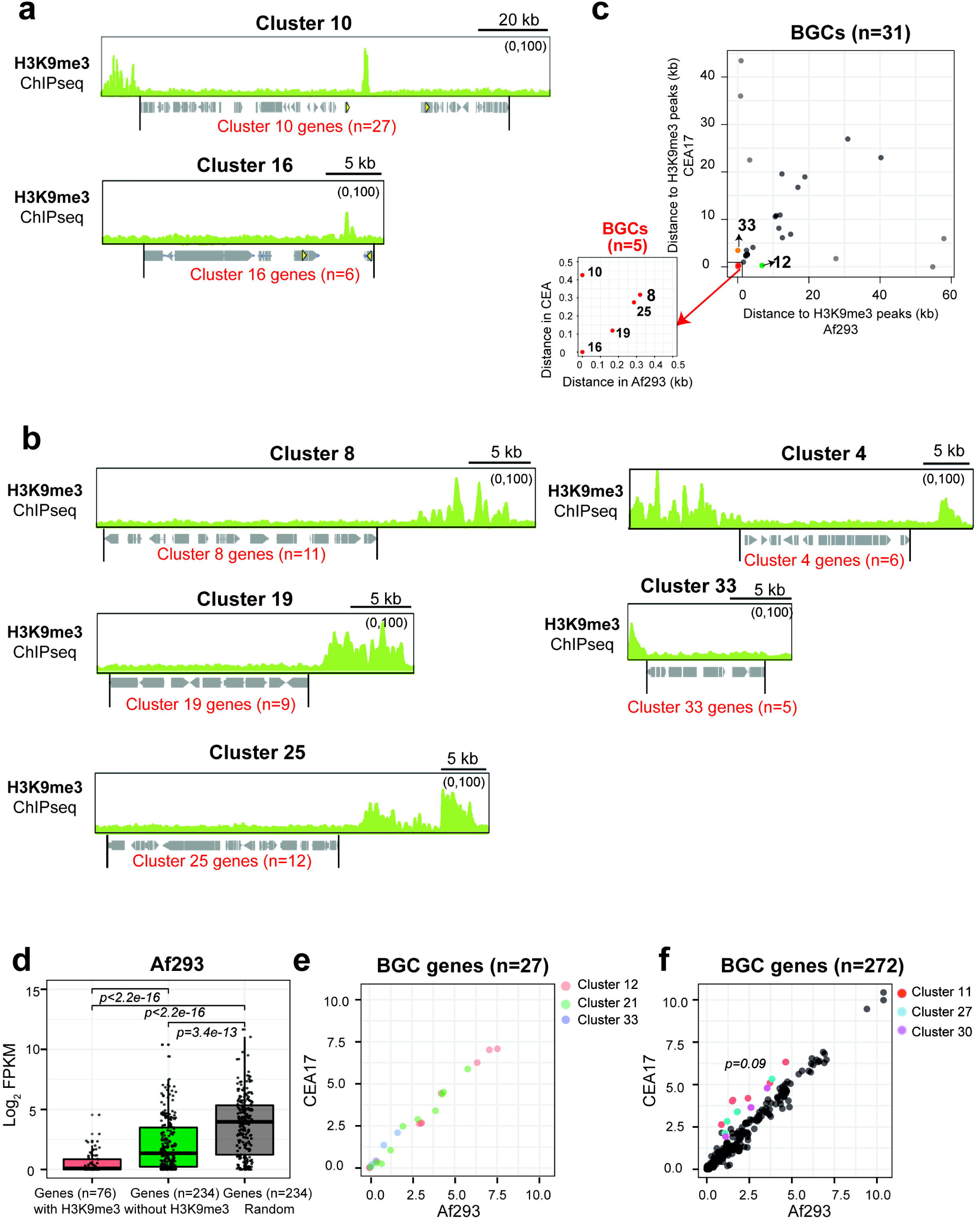
Majority of BGCs in *A. fumigatus* were not marked by H3K9me3 in Af293 and CEA17. a, Genome browser screenshots showing the selected BGCs (BGC 10 and 16) marked with H3K9me3 in Af293. b, Genome browser screenshots showing the selected BGCs marked with H3K9me3 within 4kb in Af293. c, A scatter plot showing the distance of H3K9me3 peaks to BGCs in Af293 and CEA17. d, A bar plot showing the expression of BGC genes with (red) or without (blue) H3K9me3 in Af293. Random remaining non-SM genes (grey, n = 234) was plotted as control. e, Scatter plots showing the expression of selected BGCs (12, 21, and 33) with distinct H3K9me3 modification in Af293 and CEA17. f, Scatter plots showing the expression of selected BGCs with similar H3K9me3 modification in Af293 and CEA17. Genes in BGCs 11, 27 and 33 were coloured showing higher expression in CEA17 than Af293.

It has been shown for the sterigmatocystin (ST) BGC in *Aspergillus nidulans* (Reyes-Dominguez *et al*., 2010) and lolitrem and ergot alkaloid BGCs in *Epichlo*ё *festucae* (Chujo and Scott, 2014) that H3K9me3 modification is located outside the borders of the clusters and associates with silencing of the cluster genes expression. We, therefore, extended the analysis to identify the nearest H3K9me3 peak for each SM BGC, but only found 5 additional SM BGCs with an H3K9me3 peak within 5 kb on one side of the cluster in both Af293 (BGCs 4, 8, 19, 25 and 33) (Figure 3b) and CEA17 (BGCs 8, 10, 12, 19 and 25) (Figure 3c). Hence, these results show that only seven SM BGCs in each of the isolates (out of 33 and 31 SM BGCs in Af293 and CEA17, respectively) are potentially regulated by H3K9me3 modification. Among the H3K9me3-associated clusters, five clusters (BGCs 8, 10, 16, 19 and 25) (Figure 3c) were commonly modified by H3K9me3 between the two isolates. Consistent with the heterochromatic role of H3K9me3, genes within these clusters have significantly lower expression (*i.e.,* silent) than those BGC genes without H3K9me3 modification (Figure 3d, Supplementary Figure 3d). In addition, BGC 33 was modified only in Af293 (Supplementary Figure 3e), while BGCs 12 and 21 were modified only in CEA17 (Supplementary Figure 3b, e). It is noteworthy that those BGCs were located in the same genomic loci in Af293 and CEA17 genomes, but with distinct H3K9me3 markers, suggesting that the difference is unlikely due to genomic locations. Interestingly, despite these BGCs have differential H3K9me3 modifications, the cluster genes were expressed and their expression values were highly similar between the two isolates (Figure 3e), suggesting that the observed H3K9me3 modifications did not play a major role, if any, on their gene expression. In fact, the expression values of most of the other BGC genes were highly correlated in the two isolates independent of whether their clusters were modified by H3K9me3 (Figure 3f) and were relatively much lower than the average expression level of *A. fumigatus* genes (Figure 3d, Supplementary Figure 3d). However, there are a few notable exceptions in this regard: genes of BGCs 11 [for fusarine C biosynthesis], 27 (for pyripyropene A biosynthesis) and 30 (for fumagillin biosynthesis) were expressed at noticeably higher levels in CEA17 when compared to Af293 (Figure 3f). Consistently, much higher production of pyripyropene A and fumagillin were detected in CEA17 (Supplementary Figure 3f). Taken together, the overall results indicate that, unlike what was observed for several filamentous fungal species, the majority of BGCs are not modified by H3K9me3 in *A. fumigatus*, raising the interesting question of what is the BGCs regulatory mechanism in *A. fumigatus*.

### Af293 and CEA17 genomes have distinct transposable element (TE) makeup

Since the BGCs are not the main reason for the differences in the H3K9me3 modification profiles between the two isolates, we turned to TEs that are also targets of H3K9me3 heterochromatin regulation in many other organisms (Allshire and Madhani, 2018). To systematically identify TEs, we employed the RepeatMasker program (Smith *et al*., 2015) and identified 539 and 654 TEs from the Af293 and CEA17 genomes, respectively (Supplementary Table 4). Since the CEA17 genome sequence is incomplete, the number is likely to be an underestimate. Nevertheless, it is noteworthy that the number of TEs in both genomes far exceeded the number of H3K9me3 peaks (n = 121 and 143 for Af293 and CEA17, respectively), suggesting that not all TEs were modified (and hence regulated) by H3K9me3. Alternatively, each H3K9me3 modified region may overlap with multiple TEs. Inspection on the genome browser found that some of the H3K9me3 peaks indeed overlap with several TEs (Figure 4a, b) and that strong H3K9me3 ChIP-seq signals were often found at or near TEs located in clusters (Figure 4b, indicated by arrow). On the other hand, isolated TEs tend to lack H3K9me3 modification (Figure 4b). Systematic analysis showed that around half (45%) of TEs were actually associated with H3K9me3 modification, when considering the window of 4 kb up- and down-stream of a given TE (Figure 4c).

**Figure 4.**
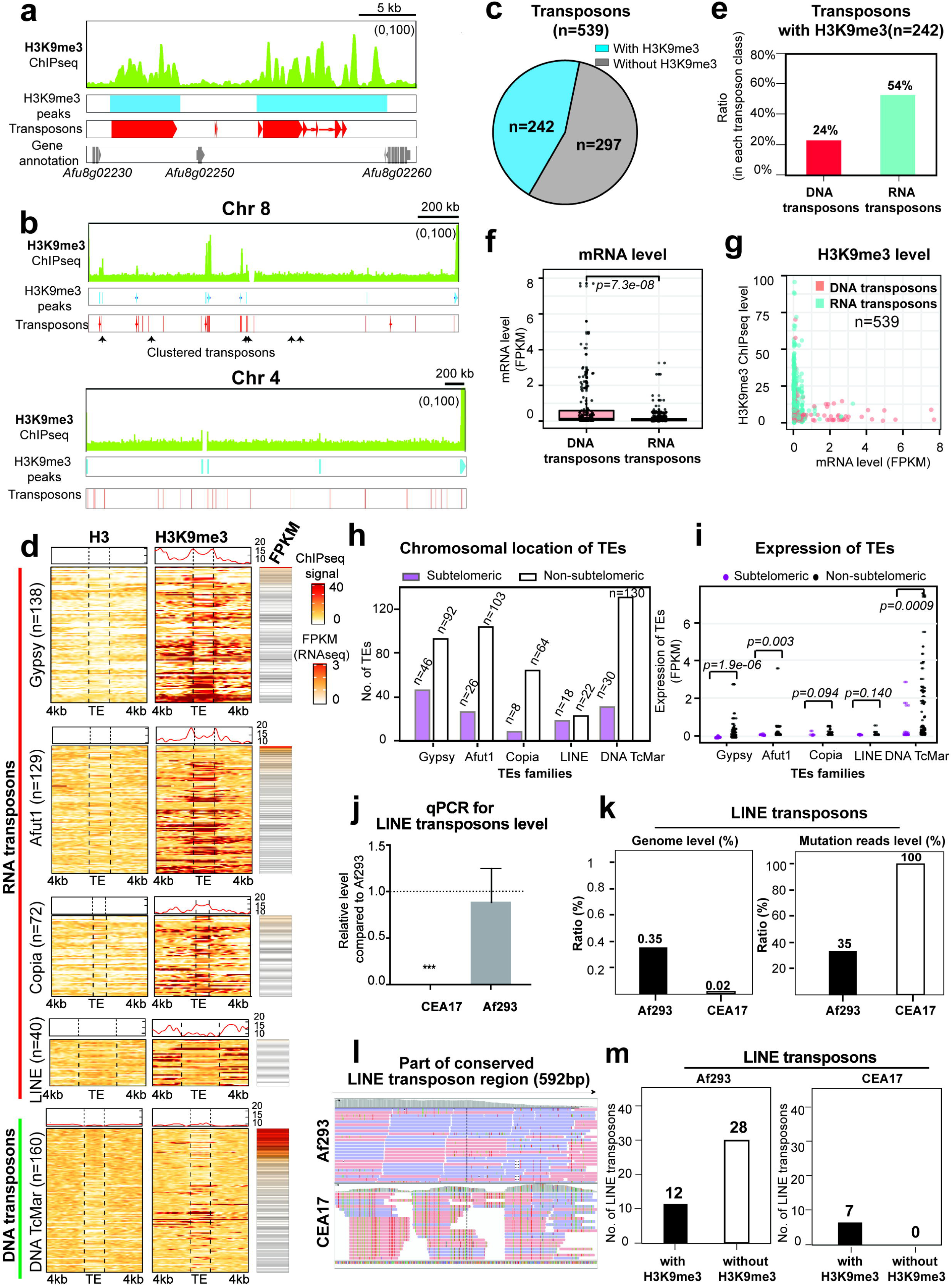
Different TEs distribution between Af293 and CEA17. a-b, Genome browser screenshots showing the distribution of H3K9me3 peaks called by MACS2 and transposons detected by RepeatMasker in Af293. c, A sector diagram showing the distribution of transposons with or without H3K9me3 genome wide in Af293. d, Heatmap plots showing the deposition levels of H3 and H3K9me3 at the Af293 TEs loci identified by RepeatMasker and the 4kb boundary regions at both sides, and the expression levels as shown by FPKM. e, A bar plot showing the number of DNA and RNA transposons bound by H3K9me3 in Af293. f, A bar plot showing the expression levels of DNA and RNA transposons in Af293. g, A scatter plot showing the relationship between expression level and H3K9me3 modification level at DNA and RNA transposons in Af293. h, A diagram plot showing the chromosomal location of different transposon families according to Repeatmasker classification in Af293. i, A scatter plot showing the expression level of different transposon families same as (h) in Af293. j, A bar plot showing the quantification of the genome level of LINE transposons in Af293 and CEA17 detected by qPCR. The qPCR was processed in two independent replicates and P value was calculated by unpaired t test. Error bar represent standard derivation and * means P value <0.05; ** means P value <0.01; *** means P value <0.005. k, A bar plot showing the genome percentage of LINE transposons in the Af293 and CEA17 genomes and their mutation reads percentage by alignment to the LINE conserved sequence in supplementary Data 5. l, A genome browser screenshot showing the alignment of Af293 and CEA17 genomic DNA to a conserved LINE transposon sequence in supplementary Data 5. The mismatched nucleotides were colorized. m, A bar plot showing the number of LINE transposons with and without H3K9me3 at the TE loci or within the boundaries in Af293 and CEA17.

To determine whether/which class of TEs are regulated by H3K9me3 modification in *A. fumigatus*, we further sub-classified the identified TE elements. Of the 539 Af293 transposon ORFs detected by Repeatmasker, 379 were classified as RNA transposons and 160 as DNA transposons (Figure 4d). Among the RNA transposons, 138 were assigned to the Gypsy group, 129 to the Afut1 group, 72 to the Copia group and 40 to the LINE group, while all the 160 DNA transposons belonged to the DNA TcMar group (Figure 4d). Interestingly, the modification pattern seemed to differ between different TEs; for example, Gypsy, Afut1 and Copia transposons were bound by H3K9me3 at their ORFs and/or 4kb flanking regions at both sides while the LINE transposons were bound by H3K9me3 mainly at the flanking regions but not within the transposon body (Figure 4d). It is also noteworthy that there was a distinct difference between DNA and RNA transposons in that more than half (54%) of RNA transposons were modified by H3K9me3, while only a quarter (24%) of DNA transposons had the histone mark (Figure 4e). Moreover, there is a significantly larger number of DNA transposons being expressed (n = 35) as compared to RNA transposons (n = 7) (Figure 4d, f), and those expressed DNA transposons were not detectably marked by H3K9me3 (Figure 4g). A similar situation was observed for CEA17, which carries 654 transposons with 476 RNA transposons (133 Gypsy, 209 Afut1, 7 LINE and 72 Copia) and 178 DNA transposons (DNA TcMar) (Supplementary Figure 4a-e). These observations indicate a preferential regulation by H3K9me3 of RNA transposons over DNA transposons.

Analysis of the chromosomal position of these elements revealed that only ∼25% (n=128) of TEs were located at subtelomeric regions and that Gypsy, Afut1, Copia and DNA TcMar TEs were predominantly located at non-subtelomeric regions. By contrast, around half (18 out of 40) of LINE TEs were at subtelomeric regions (Figure 4h). Notably, the DNA TcMar, Gypsy and Afut1 TEs located at non-sub telomeric regions had relatively higher expression levels than those situated at sub telomeric regions based on the RNAseq data, while the LINE and Copia transposons were relatively lowly expressed regardless of genomic location (Figure 4i). These results suggest a correlation between chromosomal location, H3K9me3 deposition and DNA transposons activity in Af293 (and presumably also in CEA17, whose incomplete genome assembly prevents a chromosomal-level analysis).

Notably, despite the fact that CEA17 has more TEs identified by RepeatMasker (n = 654, which is likely an underestimate due to the fragmented nature of the assembly) than Af293 (n = 539), the Af293 genome contained significantly more (*e.g.,* >5 times) LINE transposons when compared to CEA17 (Supplementary Figure 4f). For a comparison, there was less than 2-fold difference in the numbers for all the other TE classes. The absence of these Af293-unique LINE transposon regions in the CEA17 genome was further confirmed by mapping CEA17 genomic DNA sequencing data to Af293 reference genome (Supplementary Figure 4g) and by real time PCR analysis on the genomic DNA of CEA17 and Af293 (Figure 4j) using a pair of primers specific for a LINE-transposon coding ORF (Afu8g06290) that is not present in CEA17 based on BLAST analysis (Supplementary Figure 4h).

We observed a remarkable difference in the number of mutations among the LINE TEs of Af293 and CEA17 as revealed by comparisons to the consensus DNA sequence LINE TE. The LINE-transposons in the Af293 genome accumulated less mutation as compared to CEA17 (Figure 4k, l), indicating potential transposon activity from some LINE-transposons in Af293 if expressed. No difference was observed for the DNA TE Aft1 sequence (Supplementary Figure 4i) and rRNA (Supplementary Figure 4j) between the two isolates. More importantly, while all the LINE transposons in CEA17 were modified by H3K9me3 (Supplementary Figure 4a), only less than 1/3 of LINE transposons in Af293 had the histone mark (Figure 4d, m), suggesting that some of them may not be subjected to H3K9me3 silencing. Since the LINE-transposons are the most active RNA transposons and have a major impact on the genome stability (Bourque *et al*., 2018), these results imply that Af293 genome may have higher LINE-transposon activity and, hence, more unstable.

### Evidence for Af293 having a higher level of genome instability

In light of the above TE observations, we set out to assess whether Af293 had a less stable genome than CEA17. If this were true, it is expected that the Af293 strain from different laboratories may be significantly different (*i.e.,* with genetic modifications) as a result of continuous passaging over time, while the CEA17 strain accumulated relatively less changes. To assess this, we took advantage of the published RNA-seq data for the Af293 (n = 92) and CEA17 (n = 114) isolates from different laboratories to identify macro-chromosomal losses (reviewed by Priest *et al*., 2020). The rationale behind this approach is that if a strain underwent a chromosomal loss, the genes residing at the lost region would have no RNA-seq reads (or very low levels due to potential background reads from non-specific mapping). Hence, a significant continuous stretch of seemingly non-expressed (or low-expressed) genes in a given strain may indicate a chromosomal loss.

Roughly an equal number of public RNA-seq datasets were processed and analysed for the two isolates (92 and 114 datasets from 19 and 28 studies by 16 and 17 laboratories for Af293 and CEA17, respectively) (Supplementary Data 3). None of the CEA17 datasets showed noticeable signs of major chromosomal changes (Supplementary Figure 5), based on the analysis of eleven contigs including the four largest assemblies of the incomplete CEA17 genome sequence. In contrast, obvious chromosomal differences were noted in a few Af293 datasets in which some parts of a few chromosomes appeared to be missing (Figure 5a, Supplementary Figure 5), consistent with the above TE observations for instability. The potential missing regions range from 35 to 320 kb in sizes. In particular, a region of approximately 320 kb at the right end of the chromosome VIII (Chr8) appeared to be lost in the Af293 strain used in different studies (PRJNA601094, PRJNA399754, and PRJNA421149) from two different laboratories (Figure 5b, b, Supplementary Data 4). These isolates also lost other chromosomal regions (*e.g.*, the chromosomes I, V or VI) (Figure 5a), and these missing regions are not the same among the isolates. Interestingly, besides this 320 kb Chr8 region, another region of about 100 kb in length overlapping the 320 kb region was missing in another Af293 isolate from yet a different laboratory (PRJNA471263) (Figure 5a). Overall, the results suggest that the Af293 strains in different laboratories had independently acquired chromosomal losses with the Chr8 region affected in multiple independent strains, suggesting that this region is prone to chromosomal loss.

**Figure 5.**
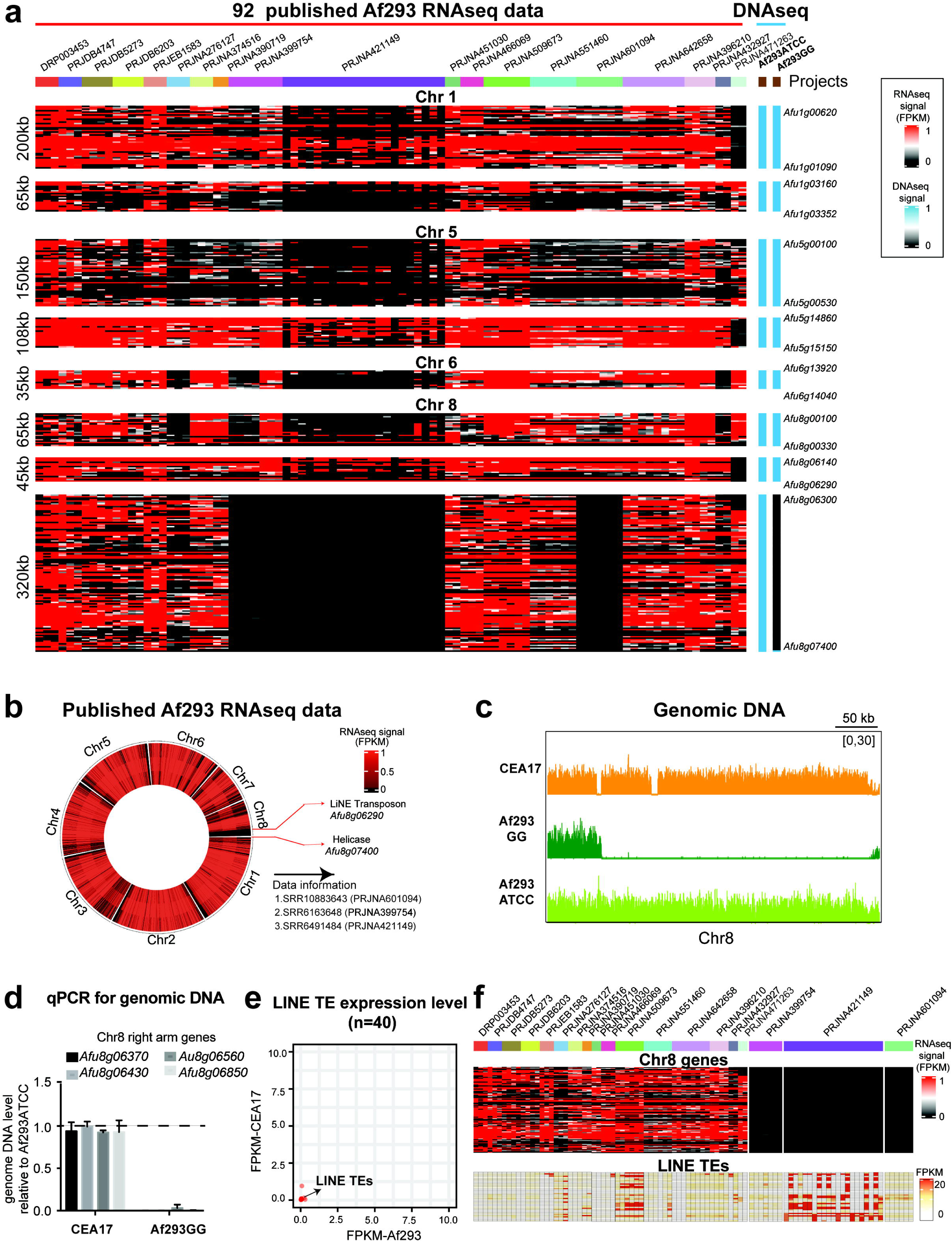
The genome instability of Af293 isolates. a, Heatmap plots showing selected chromosome regions from 92 published Af293 RNA-seq data and genomic DNA of the Af293 isolate from our laboratory (Af293GG) and the stock center (Af293ATCC). b, A circle heatmap plot showing the selected three RNAseq data sets with no reads in Chr8 right arm and the genes flanking this region. c, A genome browser screenshot showing the genomic DNA levels of Chr8 right arm of CEA17, Af293GG and Af293ATCC strains. d, A bar plot showing the levels of selected genes from the right arm of Chr8 in CEA17 and Af293GG isolate when compared to Af293ATCC. The qPCR was processed in two independent replicates and P value was calculated by unpaired t test. Error bar represent standard derivation and * means P value <0.05; ** means P value <0.01; *** means P value <0.005. e, A scatter plot showing the expression level of LINE TEs in Af293 and CEA17. f, Heatmap plots showing the LINE expression level in published 92 RNAseq data.

To further demonstrate that Af293 strains in different laboratories have genetic modifications and to confirm that the lack of RNA-seq reads in the above observation was indeed due to the loss of chromosomal regions, we sequenced the DNA of an existing Af293 culture in our laboratory (referred to as Af293GG hereafter) and a newly acquired Af293 culture from the ATCC fungal collection (referred to as Af293ATCC hereafter). Interestingly, the 320 kb of the right arm of Chr8 was also missing in our laboratory strain (Figure 5c, Supplementary Figure 6a). It is noteworthy that this is the only missing region in the Af293GG strain when compared to Af293ATCC, in contrast to the above-mentioned strains that have lost multiple regions (the right two columns in Figure 5a), indicating that Af293GG is not related to those strains (*i.e.*, not a derivative strain or a shared strain with those other laboratories). The missing right arm of Chr8 in Af293GG was further confirmed using real time PCR analysis on the genomic DNAs of the two strains and CEA17 as control (Figure 5d) and by Nanopore genome sequencing (Supplementary Text 1).

One interesting observation is that the expression of silent LINEs (Figure 5e) was elevated among some of the datasets (*e.g.*, those in PRJNA421149 project) showing the loss of the Chr8 right arm (Figure 5f), hinting that the induced LINE-transposon activity may be responsible for the chromosomal loss. However, the relationship between the Chr8 loss and these LINE TEs expressions was not strictly correlated in all datasets missing the same Chr8 region. Nevertheless, the overall results provided several lines of direct and indirect evidence that the Af293 genome is relatively unstable compared to CEA17.

### The Af293GG isolate has gained amplification of the fumitremorgin SM cluster genes and elevated production of fumitremorgin and various SMs

Genome analysis comparison revealed that the missing ∼320 kb region of Chr8 in Af293GG encompasses 109 genes including seven that encode DNA-binding transcription factors (Supplementary Table 5). Although most of these missing genes were not functionally characterized (Supplementary Table 5), GO analysis found significant enrichments for maltose metabolic process, polyamine biosynthesis and cell wall modification process (Supplementary Table 6). To determine whether the chromosomal loss in Af293GG resulted in altered gene expression, we performed transcription profiling using ChIP-seq against RNA polymerase II (Tan and Wong, 2020) for the Af293ATCC and Af293GG strains. Differentially expressed genes (DEGs) analysis identified 253 differentially transcribed genes (197 up regulated and 56 downregulated genes) in Af293GG comparing to the Af293ATCC strain (Supplementary Table 7). GO enrichment analysis showed that the down-regulated genes were enriched for plasma membrane organization, thiamine and protein transport processes (Supplementary Table 8), while the up-regulated genes showed enrichment for genes encoding stress response, carbon metabolism, reproduction, pigmentation and melanin biosynthetic process (Supplementary Figure 6b).

The enrichment of secondary metabolism in the up-regulated genes in the Af293GG prompted us to analyse the production of various known secondary metabolites by the two strains using LC-MS. Consistent with the transcription profile showing increased expression of pyripyropene A biosynthesis genes (BGC27) in Af293GG (Figure 6a), a higher level of pyripyropene A was produced by Af293GG comparatively to Af293ATCC (Figure 6b, c). In addition, a few other SMs (*e.g.*, fumitremorgin C, fumigaclavine A and gliotoxin) (Figure 6d) were also produced at significantly elevated levels by Af293GG. The detection of fumitremorgin C was unexpected, as it was reported that the Af293 isolate is incapable of producing fumitremorgin due to a point mutation (R202L) in the *ftmD* gene that compromises the methyltransferase activity by 20 folds (Kato *et al*., 2013). The DNA sequence of the *ftmD* gene in the Af293GG variant confirmed the presence of the R202L point mutation (Supplementary Figure 6c). Interestingly, in the course of inspecting the DNA sequencing data on the genome-browser, it was noticed that the Af293GG variant might have amplified a region of 55 kb on the Chr8 left arm adjacent to a LINE-TE Afu8g00310 (Figure 6e). This was confirmed by Nanopore genome sequencing (Supplementary Text 1). The amplification was further corroborated by quantitative PCR using primer pairs specific to three different loci within the region (Supplementary Figure 6d). The quantitative PCR result showed a 2-fold increase in the DNA level, suggesting a duplication of this region. The duplication encompasses 20 genes with eleven being uncharacterized and nine belonging to the fumitremorgin A-C biosynthetic cluster (Supplementary Table 9). The amplification of the cluster was further confirmed by quantitative PCR on three of the cluster genes [*fmtC* (Afu8g00190), *fmtD* (Afu8g00200) and *fmtE* (Afu8g00220)] (Figure 6f) and might have consequently increased the expression of these cluster genes including the *ftmD^R202L^* mutant gene, thereby increasing fumitremorgins production in Af293GG. Indeed, RT-qPCR detected 5-to-10-fold higher mRNA levels on three of the cluster genes (*ftmC*, *ftmD* and *ftmE*) in Af293GG when compared to Af293ATCC, and their levels were even higher than those in CEA17 (Figure 6g). However, there was no difference in H3K9me3 modification between Af293GG and Af293ATCC at the fumitremorgin cluster (Figure 6h) as well as all other SM clusters (Figure 6i), which is in agreement with the lack of association between BGCs regulation and H3K9me3 modification described above., which is in agreement with the lack of association between SM gene silencing and H3K9me3 modification described above.

**Figure 6.**
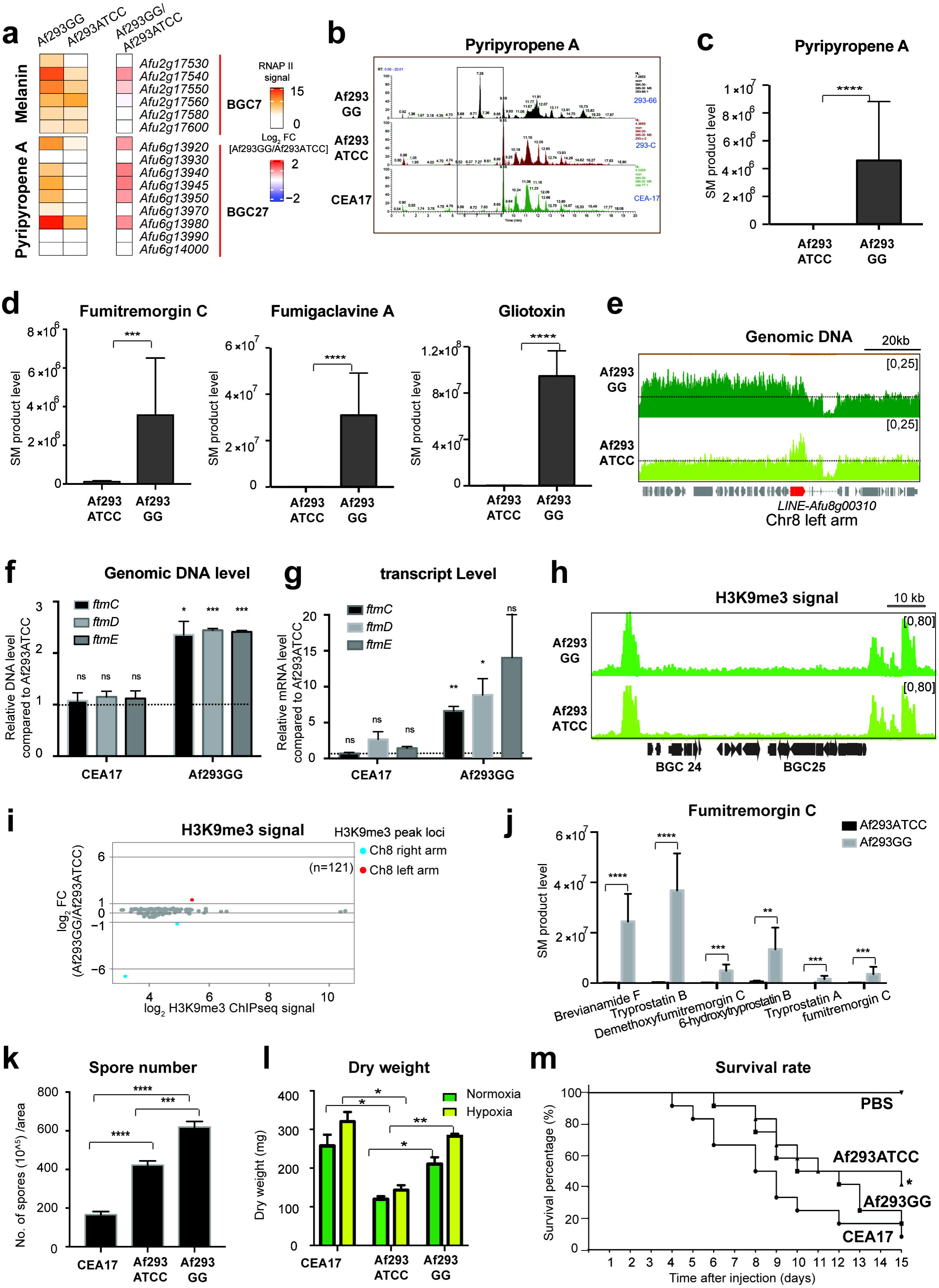
The evolved Af293GG isolate induced production of many SM products and gained amplification of chr8 left arm, better growth fitness and higher virulence. a, Box plots showing the expression level detected by RNAP II ChIPseq of melanin and pyripyropene A biosynthetic genes in Af293GG and Af293ATCC. b, LC/MS profiles showing the peaks representing pyripyropene A products in Af293GG, Af293ATCC and CEA17. c-d, Bar plots representing the quantified production of SMs (c) pyripyropene A as shown in (b) and fumitremorgin C, fumigaclavine A and gliotoxin (d) in Af293ATCC and Af293GG. The SMs levels were calculated as peak area, and P values were calculated by F test to compare variations. Error bar represent standard derivation and * means P value <0.05; ** means P value <0.01; *** means P value <0.001, **** means P value <0.0001. e, A genome browser screenshot showing the genomic DNA level of Chr8 left arm of CEA17, Af293GG and Af293ATCC. The transposon ORF Afu8g00310 is marked in red. f, A bar plot showing the relative genomic DNA level detected by qPCR of selected coding (*fmtC*, *ftmD* and *ftmE)* regions of Chr8 left arm in CEA17 and Af293GG genomes when compared to Af293ATCC. The qPCRs were processed in two independent replicates and P values were calculated by unpaired t test. Error bar represent standard derivation and * means P value <0.05; ** means P value <0.01; *** means P value <0.005. g, A bar plot showing the qPCR relative quantification of the expression levels of *fmtC-E* genes in CEA17 and Af293GG isolates when compared to Af293ATCC. The qPCRs were processed in two independent replicates and P values were calculated by unpaired t test. Error bar represent standard derivation and * means P value <0.05; ** means P value <0.01; *** means P value <0.005. h, A genome browser screenshot showing the H3K9me3 deposition at the boundaries of BGC 24 and BGC 25 in Af293ATCC and Af293GG. i, A MA plot representing the relative H3K9me3 ChIP-seq signal in Af293GG when compared to Af293ATCC. The red dot is the peak located at the Chr8 left arm, and the two blue dots are the peaks located at the Chr8 right arm. j, A bar plot showing the quantified production of fumitremorgin C and its intermediates in Af293GG and Af293ATCC as shown in Supplementary Figure 7b, Error bar represent standard derivation and * means P value <0.05; ** means P value <0.01; *** means P value <0.001, **** means P value <0.0001 as shown by F test to compare variations. k, A bar plot showing the number of spores per area produced by CEA17, Af293ATCC and Af293GG strains. l, A histogram plot showing the dry weight of CEA17, Af293ATCC and Af293GG biomass when the strains were grown in static liquid cultures exposed to normoxia (∼21% O_2_) and hypoxia (0.9% O_2_) conditions for 96 h. m, Survival curves (n=12/strain) of *G. mellonella* infected with CEA17, Af293GG and Af293ATCC spores. Phosphate buffered saline (PBS) without conidia was used as a negative control. Indicated P-values are based on the Log-rank, Mantel-Cox and Gehan-Breslow-Wilcoxon tests. * means P value <0.05. -ATCC isolates. Phosphate buffered saline (PBS) without conidia was used as a negative control. Indicated P-values are based on the Log-rank, Mantel-Cox and Gehan-Breslow-Wilcoxon tests. * means P value <0.05.

Fumitremorgins are indole alkaloids, whose biosynthesis begins by condensation of the two amino acids L-tryptophan and L-proline to brevianamide F, catalyzed by FtmA. Brevianamide F is then prenylated by FtmPT1/FtmB, resulting in the formation of tryprostatin B. FtmE is responsible for the conversion of tryprostatin B to demethoxyfumitremorgin C and tryprostatin A to fumitremorgin C, while the conversion of tryprostatin B to 6-hydroxytryprostatin B and 6-hydroxytriprostatin B to tryprostatin A are catalyzed by FtmC and FtmD, respectively. The subsequent reactions result in the formation of 12,13-dihydroxyfumitremorgin C, fumitremorgin B, verruculogen and fumitremorgin A (Shu-Ming Li, 2010). Consistent with the biosynthetic pathway, LC-MS analysis revealed that all the intermediates in the fumitremorgin biosynthetic pathway (Supplementary Figure 7a) were produced in higher amounts by Af293GG (Figure 6j, Supplementary Figure 7b). Moreover, there was a clear distinction in the levels of intermediate metabolites before and after the biosynthesis step mediated by FtmD^R202L^: the intermediates brevianamide F, tryprostatin B and 6-hydroxytriprostatin B, which are produced before the step limited by the mutated enzyme were accumulated at higher levels comparing to the intermediates tryprostatin A and fumitremorgin C, which are produced in the subsequent steps (Figure 6j). This observation is consistent with the reduced methyltransferase activity of FtmD^R202L^, which imposes a rate-limiting step in fumitremorgin biosynthesis. The increased *fmtD^R202L^* expression, presumably together with elevated levels of precursor intermediate metabolites, could augment the compromised methyltransferase activity of FtmD^R202L^ for fumitremorgin production. Alternatively, another methyltransferase could be acting in this pathway. Taken together, these results demonstrated that our evolved Af293GG strain has acquired enhanced abilities for SM production as a result of major chromosomal changes (*i.e.,* amplification and loss).

### The Af293GG isolate has increased asexual spore production, growth and virulence

Growth analysis showed that Af293GG and Af293ATCC as well as CEA17 have slightly different colony morphology (Supplementary Figure 8a) and colony colour, which may indicate a variation in spore production. Indeed, the three strains produced different numbers of asexual spores under the same conditions with Af293GG having the most (Figure 6k). Moreover, Af293GG also generated higher biomass than Af293ATCC both in normoxia and hypoxia growth conditions at levels similar to CEA17 (Figure 6l). Given the positive effects of hypoxia fitness (Kowalski *et al*., 2016) and mycotoxin production (*e.g.,* gliotoxin) towards *A. fumigatus* virulence, we next asked whether the evolved Af293GG isolate also has increased virulence. We infected *Galleria mellonella* larvae with Af293, Af293GG and CEA17 conidia and monitored the infection attibutes such as survival rate, worm movement and cuticle melanisation for 15 days. The larvae infected by CEA17 conidia began to die on the 4^th^ day, while the larvae infected by Af293 and Af293GG conidia started dying on the 6^th^ day (Figure 6m). After 11 days, the larvae infected by Af293GG conidia were more melanized and less active than the ones infected by Af293 (Supplementary Movie 1). On the 15^th^ day, 8%, 40% and 16% of the larvae infected by CEA17, Af293 and Af293GG strains, respectively, were alive (Figure 6m) and the median survival day for each strain were 8.5, 13 and 11 days, respectively (Supplementary Figure 8b). Although all three strains were able to cause death eventually, the observed median survival rate, movement and melanisation levels of infected *G. mellonella* collectively indicate subtle but significant differences between our lab evolved Af293GG strain, the standard Af293ATCC and CEA17 isolates.

## Discussion

In this work, we performed the first *A. fumigatus* chromatin profiling for two histone modifications, H3K4me3 and H3K9me3, to identify the similarities and differences between the two most commonly studied clinical isolates, Af293 and CEA17. We showed that the transcriptional activity positively correlates with promoter nucleosome depletion and H3K4me3 modification at 5’ end of transcribed genes, which is consistent to what has been observed in other eukaryotes. H3K4me3 recruits proteins involved in RNAP II transcription and preferentially marks highly expressed, environmentally insensitive (*i.e.*, housekeeping) genes (Murray *et al*., 2019). Our results reinforce this idea since in both strains of *A. fumigatus* highly expressed genes are marked by H3K4me3 participating on primary metabolism and cellular component organization and biosynthesis, while unmarked genes that have low expression are involved in secondary metabolism and cell wall remodelling processes. Notably, a specific gene cluster codifying several transporters were not marked by this modification, which is consistent with previous findings that H3K4me3 is not instructive for the expression of inducible genes (Howe *et al*., 2017). One explanation for this is that the transcription of those genes with defined function does not require H3K4me3 (Pérez-Lluch *et al*., 2015). Another possible explanation is related to the fact that ChIP-seq experiments capture, through formaldehyde crosslinking, events that occur at the time of cell harvest. Therefore, those transporter genes may be transcribed earlier before the actual experiment (*i.e.,* during spore germination or the initial growth stage) but their mRNAs are stable enough until the point of experiment for detection by RNAseq, whereas H3K4me3 was removed soon after transcription ceased. Accordingly, a transcriptome analysis of germinated spores (Lamarre *et al*., 2008) supports the second hypothesis, as it showed an induction of transporter genes (*e.g.* ABC drug efflux pumps (*abcB* and *abcD*) and MFS (*msfB* and *msfC*) at early germination stages (30 min) that decreased at later time points (90 min).

The histone modification H3K9me3, which is usually associated with heterochromatin (Allshire and Madhani, 2018), was also systematically investigated. Here we demonstrated that not only most of *A. fumigatus* BGCs are not marked by H3K9me3 but also that the modification presence does not always mean BGC repression. Of note, we detected more silent BGCs among the ones nearby the H3K9me3 loci in Af293 and CEA17, suggesting that these BGCs could be regulated by H3K9me3. In fact, a variable relationship has been shown between these two elements in filamentous fungi. For instance, while the H3K9me3 removal led to an increased expression of several BGCs in *A. nidulans* (Reyes-Dominguez *et al*., 2010), only a partial de-repression was achieved in the plant endophyte *Epichloë festucae* (Chujo and Scott, 2014), and no effect was observed in the plant pathogen *Zymoseptoria tritici* (Moller *et al*., 2019). The previous and our results together suggest that H3K9me3 is less important for regulation of most SM clusters than previously thought.

The Af293 genome is 1.4% smaller than CEA17 genome (Fedorova *et al*., 2008), but contains more transposon related ORFs according to RepeatMasker software. Here we demonstrated a higher number of retrotransposons than DNA transposons in both isolates, and the retrotransposons were more marked by H3K9me3. A similar situation was previously detected in *C. elegans* embryos, where 58.5% of retrotransposons and 17.7% of DNA transposons carried the H3K9me3 modification (Zeller *et al*., 2016). In fact, the DNA transposons were more active in our RNA-seq, especially those at non-subtelomeric regions, suggesting that both chromosomal location and H3K9me3 deposition regulate the transposons expression and hence activity in *A. fumigatus.* The relationship between the histone PTM and transposons is consistent with other eukaryotes, such as *Z. tritici* (Moller *et al*., 2019) and *C. elegans* (Zeller *et al*., 2016), where loss of H3K9me3 increases the expression of both TE classes. A closer look into the different transposon families revealed that Af293 genome contains several LINE-transposons, which were absent in CEA17. This finding agrees with the previous detection of 24 LINE-like elements (LLEs) in Af293 in contrast to only 4 heavily mutated LLEs in CEA17 (Huber and Bignell, 2014). Here, we described that half of Af293 LINE-transposons were located at subtelomeric regions and contained H3K9me3 modification only at their boundaries, while the other retrotransposon families (*e.g.,* Gypsy, Afut1 and Copia) were mostly localized at internal chromosomal regions and bound by H3K9me3 along their entire length, implying more activity of LINE-transposons when compared to other families. These results are also consistent with recent findings in the fungal plant pathogen *Verticillium dahliae,* where individual transposon families have distinct epigenetic and compactation profiles (Cook *et al*., 2020). In this species, the younger transposons display lower level of nucleotide divergence from the consensus sequence and are more transcriptionally active, as opposed to older transposons, which are more mutated and less active (Faino *et al*., 2016). The same situation was observed for Af293 where the larger number of Af293 retrotransposons results from recent insertions (*i.e.,* younger transposons) (Huber and Bignell, 2014). We observed that LINE TEs have increased numbers and less mutations in Af293 compared to CEA17. These results together with the data provided heresuggest that Af293 LINE-transposons could be activated under proper environmental stimuli. In particular, several stresses, such as high temperature, have been shown to trigger transposons activity (Zeller *et al*., 2016; Miousse *et al*. 2015), which has been associated with chromosome and genome instability in different organisms and cell lines (Moller *et al*., 2019, Zeller *et al*., 2016; Kondo *et al*., 2008; Peters *et al*., 2001; Peng *et al*, 2009). Overall, our study indicates that the absence of H3K9me3 in Af293 LINE-transposons ORFs potentially contribute to genomic and chromosomic rearrangement events, such as deletions, duplications and chromosome breakages.

The search for a possible relationship between a change in gene expression and chromosomal alterations in public Af293 and CEA17 RNAseq datasets returned distinct levels of genomic alterations between the two isolates, indicating that different Af293 genomes are more variable when compared to CEA17 datasets. The possible occurrence of a similar large deletion of approximately 320 kb at the right arm of Chr8 was observed in three Af293 isolates from two distinct laboratories following this approach while no obvious changes were detected in CEA17-derived isolates. Interestingly, the transcriptional profiling for the Af293 isolates used in these three studies was performed in mice and cell culture infection experiments. It is tempting tospeculate that stressing infection-associated factors, such as fever and/or immunological attacks could have contributed to the observed chromosomal changes. In *Candida glabrata*, chromosomal rearrangements are largely detected in clinical isolates acting as a virulence mechanism and driving their adaptation to different host niche environments (Poláková *et al*. 2009). As transposons are enriched near structural rearrangement breakpoints and are intrinsically linked to genome evolution (Schrader *et al*., 2014; Faino *et al*., 2016), the fact that the gene adjacent to the gap region was a LINE-transposon led us to hypothesize that this event could be the result of transposon activity on the original Af293 genome. Interestingly, this transposon was highly upregulated in one RNA-seq public project with Chr8 loss, suggesting that this event might be triggered in a particular circumstance when transposons tend to be more active (Fouché *et al*., 2019). Even more remarkable finding was the detection of an identical Chr8 right arm loss event in our own Af293 isolate, named Af293GG, which was independent of the two others above mentioned, implying that this region is a hotspot for rearrangement in Af923. The closer inspection of the Af293GG genome showed a DNA amplification region on the left arm of the same chromosome, which was also flanked by a LINE-transposon. Recent work in *Z. tritici* showed that methyltransferase mutants unable to trimethylate the histone H3 lysine 9 (H3k9me3) or lysine 27 (H3k27me3) resulted in a higher degree of genome instability (Moller *et al*., 2019). After 50 transfers of mitotically growing H3K9me3-devoid cells, they detected shortening of chromosome 6, absence of chromosome 20 and duplication of a long region on chromosome 1, which re-localized in two new independent chromosomes (Moller *et al*., 2019). Similar to what was detected here in the Chr8 of Af293GG, the breakpoint of the duplicated region in *Z. tritici* coincided with a transposon-rich region normally associated with H3K9me3 in the wild type (Moller *et al*., 2019). These observations reinforce the hypothesis that links transposons abundance and expression with increased frequency of chromosomal rearrangements in Af293.

The amplified region in the left arm of Chr8 contais the BGC for fumitremorgin, while the lost right arm portion contained six transcription factors encoding genes and genes that codify proteins involved in cell wall metabolism. This led us to speculate that the amplification, the loss or both events could render an advantage to Af293GG over Af293. Significantly, an increased production of several SMs, including fumitremorgin C, fumigaclavine A, pyripyropene A and gliotoxin was detected in Af293GG. FtmD, in the fumitremorgin cluster, encodes an O-methyltransferase that catalyzes the conversion of 6-hydroxytryprostatin B into tryprostatin A (Kato *et al*., 2013). The Af293 has a R202L mutation that blocks fumitremorgin B synthesis in this strain (Kato *et al*., 2013). Since the same mutation is detected in Af293GG *fmtD* gene, the much higher and detectable production of fumitremorgin C and its intermediates in Af293GG strain could be due to the augmented FmtD residual activity caused by its overexpression or another *O*-methyltransferase that functions at this biosynthetic step. The increased production of other secondary metabolites inAf293GG strain (*e.g.*, fumigaclavine A, pyripyropene A and gliotoxin) could be due to multiple mechanisms including specific regulation by one of the TFs lost from Chr8 right arm and/or the metabolic imbalance caused by the fumitremorgin overexpression. Although most BGCs contain specific transcription factors involved in the transcriptional regulation of determined SM clusters, an increasing number of evidences have demonstrated cross-regulation between the different BGCs (Perrin *et al*. 2007, Wiemann *et al*. 2013, Wiemann *et al*. 2014, Doyle *et al*., 2017). For instance, an iron dependent network composed by SreA and HapX induces the hexadehydroasthechrome (HAS) BGC under iron excess (Wiemann *et al*. 2014). The *hasA* overexpression, in turn, results in downregulation of several genes involved in pyripyropene, fumitremorgin and gliotoxin biosynthesis (Wiemann *et al*. 2014). In fact, gliotoxin and fumitremorgin regulation mechanisms are probably related, as the disruption of dithiol gliotoxin *bis*-thiomethylation by *gmtA* deletion results in decreased gliotoxin, tryprostatin B and fumitremorgin in *A. fumigatus* (Doyle *et al*., 2017). Inversely, here we show gliotoxin production as a result of the fumitremorgin BGC upregulation, reinforcing the idea that specific fungal BGCs are interconnected and controlled by diverse regulatory mechanisms.

Overall, our work emphasizes the importance of epigenetic modifications in *A. fumigatus* strain heterogeneity as evolutionary drivers of altered gene expression and genome stability. The unstable Af293 genome was characterized compared to CEA17 and a new strain Af293GG with Chr8 rearrangement was isolated in the laboratory. When compared to the parental strain Af293GG gained better growth fitness in both normoxia and hypoxia conditions, had higher production of several SMs and increased virulence in larvae infection, strongly indicating that the Chr8 rearrangements could be a strategy to ensure *A. fumigatus* better fitness.

## Acknowledgments

We thank Fundação de Amparo à Pesquisa do Estado de São Paulo (FAPESP) grants 2016/07870-9 (GHG), 2018/14821-0 (ACC), 2016/12948-7 (PAC), 2018/00715-3 (CV), 2019/06359-7 (TPF) 2019/15675-0 (RSR) and Conselho Nacional de Desenvolvimento Científico e Tecnológico (CNPq) (GHG), both from Brazil for financial support. We also acknowledge the support from the Research Services and Knowledge Transfer Office (project reference number: MYRG2018-00017-FHS and MYRG2019-00099-FHS) of the University of Macau, The Science and Technology Development Fund, Macao S.A.R (FDCT) (project reference number: 0106/2020/A) and the Collaborative Research Fund Equipment Grant (C5012-15E) from the Research Grant Council, Hong Kong Government. This work was performed in part at the High-Performance Computing Cluster (HPCC), which is supported by the Information and Communication Technology Office (ICTO) of the University of Macau. We thank Jacky Chan for technical supports on the HPC. We also thank Dr. Alessia Buscaino for reading the manuscript draft and for her comments and suggestions.

---

**Supplementary Figure 1. The similar nucleosome depletion and H3K4me3 modification in two *A. fumigatus* isolates Af293 and CEA17.** a, Line plots showing the H3 and H3K4me3 deposition within 1kb of TSS region of genes genome wide in CEA17 when the data was mapped to Af293 and CEA17 (A1163) reference genome, respectively. Gene order was ranked by mRNA level from high to low. The pink shade was plotted as shown the mapping was performed to CEA17 genome reference. b, Scatter plots showing the correlation of mRNA level (upper panel) and H3K4me3 deposition level (bottom panel) in Af293 and CEA17. c, Genome browser screenshots showing the mRNA and H3K4me3 level at selected genes in CEA17. H3 was used as control. d, Gene Ontology analysis of gene sets in cluster 1-4 as shown in Figure 1e, f.

**Supplementary Figure 2. The H3K9me3 modification profile in CEA17**. a, A heatmap plot showing the H3K9me3 levels in Af293 genome. Each bin represents 200bp; 100kb nearby chromosome end, 20kb pericentromere, and randomly selected 100kb regions were plotted for each chromosome. b, A genome browser screenshot showing the H3K9me3 deposition profile throughout Af293 genome reference chromosome II (upper panel) or CEA17 (A1163) genome reference Scf_000002 in Af293 and CEA17. H3 was used as control. The pink shade was plotted as shown the mapping was performed to CEA17 genome reference. c-d, Bar plots showing the distribution of selected genes located in subtelomere (c, n = 1578) and pericentromere (d, n = 70) in clusters as shown in Figure 1e, f. The colour of each bar was marked as cluster information accordingly. e, Genome browser screenshots showing the H3K9me3 and H3K4me3 profile in pericentromere loci throughout Af293 genome reference. Centromere was indicated as blue box.

**Supplementary Figure 3. Majority of BGCs in *A. fumigatus* were not marked by H3K9me3 in Af293 and CEA17.** a, A scheme diagram showing the inverted distribution of BGCS 9-16 in Af293 Chr3 and CEA17 Scf-03. The pink shade was plotted as shown the mapping was performed to CEA17 genome reference. b, Genome browser screenshots showing the selected BGCs [BGC16 and 21 marked with H3K9me3 in CEA17. c, Bar plots showing the H3K9me3 peak span ratio along with the length of its bound BGCs in Af293 and CEA17. d, A bar plot showing the expression of BGC genes with (red) or without (blue) H3K9me3 in CEA17. Random remaining non-BGC genes (grey, n= 213) were plotted as control. e, Genome browser screenshots showing the selected BGCs (BGC12, 21 and 33) with different H3K9me3 marker in Af293 and CEA17. The pink shade was plotted as shown the mapping was performed to CEA17 genome reference. f, Bar plots showing the quantified production of SMs pyripyropene A and fumagillin in Af293 and CEA17. The SMs levels were calculated as peak area, and P values were calculated by F test to compare variations. Error bar represent standard derivation and * means P value <0.05; ** means P value <0.01; *** means P value <0.001, **** means P value <0.0001.

**Supplementary Figure 4. Different TEs distribution between Af293 and CEA17.** a, Heatmap plots showing the H3 and H3K9me3 deposition at CEA17 transposon loci identified by Repeatmasker and at their boundary regions (+/− 4kb) and their expression values (in FPKM). b, A sector diagram showing the distribution of transposons with or without H3K9me3 genome wide in CEA17. c, A bar plot showing the number of DNA and RNA transposons bound by H3K9me3 in CEA17. d, A bar plot showing the expression level of DNA and RNA transposons in CEA17. e, A scatter plot showing the relationship between expression level and H3K9me3 modification level at DNA and RNA transposons in CEA17. f, A bar plot showing the distribution in families of Af293 and CEA17 TEs according to Repeatmasker classification. g, Heatmap plots showing the LINE transposon level in Af293 and CEA17. h, A bar plot showing the number of blast hits for LINE transposon in Af293 and CEA17. i-j, Bar plot showing the genome percentage and mutation reads percentage of (i) Aft1 transposons and (j) 18S rRNAs in Af293 and CEA17.

**Supplementary Figure 5. Different genome stability inferred from comparable published RNAseq data of Af293 and CEA17.** Heatmaps showing the mapped reads in genes of all chromosomes in Af293 data (n=92) and CEA17 data sets (n=114) from 19 and 28 studies, respectively.

**Supplementary Figure 6. The different genome arrangement of evolved Af293GG isolate compared to Af293ATCC isolate.** a, A heatmap plot showing the genome density of Chr8 in CEA17, Af293GG and Af293ATCC. b, A scheme diagram plot showing the Gene Ontology result of Af293GG up and down-expressed genes measured by RNAP II ChIPseq. c, A genome browser screenshot showing the SNP mutation of R202L in Af293GG and Af293ATCC isolates compared to CEA17. d, A bar plot showing the relative genomic DNA level detected by qPCR of selected non-coding regions of Chr8 left arm in CEA17 and Af293GG genomes when compared to Af293ATCC. The qPCRs were processed in two independent replicates and P values were calculated by unpaired t test. Error bar represent standard derivation and * means P value <0.05; ** means P value <0.01; *** means P value <0.005.

**Supplementary Figure 7. The induced fumitremorgin production by evolved Af293GG isolate compared to Af293ATCC isolate.** a, A scheme diagram showing the (top panel) genome distribution of fumitremorgin cluster (BGC 29) genes and (bottom panel) pathway of fumitremorgin A-C biosynthesis. b, The LC/MS profile of compounds 1-6 in the supernatants of in Af293GG, Af293 and CEA17 strains as shown in (a) and Figure 6j.

**Supplementary Figure 8. The evolved Af293GG isolate produced more spores and had higher virulence.** a, Photos showing the different colony morphologies of CEA17, Af293ATCC and Af293GG isolates. b, A histogram plot showing the median survival day of the larvae as shown in Figure 6m.

**Supplementary Table 1.** The mapping records of Af293 and CEA17 ChIP-seq data.

**Supplementary Table 2.** The putative centromere loci of Af293.

**Supplementary Table 3.** The annotated BGC genes information in Af293 and CEA17.

**Supplementary Table 4.** The TE elements identified from RepeatMasker program in Af293 and CEA17.

**Supplementary Table 5.** Genes in the right arm of Chr8 which were lost in Af293GG isolate.

**Supplementary Table 6.** GO enrichment analysis of the Af293GG lost Chr8 genes.

**Supplementary Table 7.** Differential expressed genes in Af293GG and Af293ATCC isolate detected by RNAP II ChIP-seq.

**Supplementary Table 8.** GO enrichment analysis of the Af293GG induced and repressed genes.

**Supplementary Table 9.** Genes in the left arm of Chr8 which had their copy number amplified.

**Supplementary Table 10.** Antibodies used in this project.

**Supplementary Table 11.** Oligos used in this project.

**Supplementary Data 1.** Gene Ontology analysis results from FungiDB of genes in clusters 1-4 as shown Supplementary Figure 1d.

**Supplementary Data 2.** H3K9me3 peaks detected in Af293 and CEA17 genomes.

**Supplementary Data 3.** Published 92 and 114 RNA-seq data sets for Af293 and CEA17, respectively, that were analysed in this work.

**Supplementary Data 4.** Published RNA-seq data with Chr8 right arm partial loss that were analysed in this work.

**Supplementary Data 5.** Conserved 18S rRNA, LINE transposon and Aft1 transposon sequence used for analysis in this work.

**Supplementary Text 1.** Nanopore genome sequencing results of Af293ATCC and Af293GG strains.

**Supplementary Movie 1.** A movie showing the G. melonela movement and myelinization after 3 days of infection with Af293ATCC (first petri dish) and Af293GG (second petri dish).

## Methods

### Strains and culture conditions

Two *A. fumigatus* clinical isolates, Af293 (named later as Af293ATCC) and CEA17, and one spontaneous Af293 variant (named later as Af293GG) were used in this project. Conidia were harvest after 2 days of incubation at 37 °C on complete medium agar plate. Mycelia samples were obtained by inoculating 5×10^^7^ aforementioned conidia in ANM medium with sodium nitrate as nitrogen source, cultivated in an orbital shaking incubator at the speed of 220 rpm at 37 °C for 16 hours. For the hypoxia assay, the conidia were inoculated in flasks (125 ml Erlenmeyer’s) containing 30 ml of liquid ANM medium and incubated without agitation for 96 hours at 0.9% O_2_. The flasks incubated under normal atmospheric conditions (normoxia) were used as control.

### Chromatin preparation

For DNA crosslinking, formaldehyde was added to the cultures at 1% final concentration and gently shook for 20mins at room temperature. Next, a final concentration of 0.5M glycine was added to further incubation of 10 mins. The mycelia were harvest by filtering and washed with cold water, followed by pressing in dry paper. The above crosslinked mycelia cells were snap-frozen in liquid nitrogen and frozen-dried for 3-4 hours before lysis. The cell lysis was processed by 6 times beating for 3 minutes with ∼100 μl volume of silica beads using Bullet Blender (Next Advance) with at least 3 minutes of cooling in between each cycle. Chromatins were extracted as described before (Fan, Lamarre-Vincent, Wang, & Struhl, 2008) and sonicated using the Qsonica Q800R at 100% amplitude with 10 seconds ON and 15 seconds OFF cycles for a total sonication time of 30 minutes. Chromatin concentration and size (100-500bp) were checked on 2% agarose gel, and the prepared chromatins were stored at −80 L until use.

### Chromatin Immuno-precipitation and sequencing library preparation

Immuno-precipitation was carried out as previously described (Wong and Struhl, 2011) using antibodies listed in supplementary Table 10. Immuno-precipitated materials were purified using QIAGEN PCR cleanup kit (cat no. 28106) for library preparation as described previously (Wong *et al*., 2013) with the exception of the end-repair step using NEBNext® Ultra II End Repair/dA-Tailing module (NEB, cat. no. E7546L) accordingly to manufacturer’s protocol. One ng of chromatin DNA was used for library preparation as input DNA control. Libraries were checked and quantified using DNA High Sensitivity Bioanalyser assay (Agilent, cat. no. XF06BK50), mixed in equal molar ratio and sequenced using the Illumina HiSeq2500 platform at the Genomics and Single Cells Analysis Core facility at the University of Macau.

### Genomic DNA extraction and real-time quantitative PCR (qPCR)

Genomic DNA of *A. fumigatus* was extracted from frozen mycelia as described (B.Lee & W.Taylor, 1990). One ng of DNA was used for real-time qPCR analysis using Premix Ex Taq DNA polymerase (Takara, cat. no. RR039W) on ABI Fast 7500 (Applied Biosystems®) Real-Time PCR machine and the oligos were listed in Supplement Table 11. The ΔΔCt method was used for quantification using tubulin gene *tubA* as internal reference.

### DNA sequencing using Oxford Nanopore

For long read DNA sequencing, 4 μg of genomic DNA was used to construct sequencing library using the Ligation Sequencing Kit (SQK-LSK109, Oxford Nanopore) according to the manufacture protocol. Flongle flowcells priming and loading steps were then carried according to the steps described in the manufacturer’s protocol. Sequencing runs lasted for up to 24h and were performed in MinION model Mk1B (Oxford Nanopore). FASTQ files were generated by Guppy basecaller version 4.0.9 (Oxford Nanopore) in the fast basecalling setting during sequencing. Reads that passed the default quality filter (average score ≥ 7) had barcode sequences removed by guppy_barcoder command line tool.

The two strains of *A. fumigatus* were assembled using different methodologies. First, Illumina reads were trimmed using trimmomatic v0.38 (Bolger *et al*., 2014) and long reads were trimmed using a combination of porechop v.0.2.4 (Wick *et al*., 2017) and Nanofilt 2.5.0 (De Coster *et al*., 2018). Long reads were then corrected using CANU v1.8 (Koren *et al*., 2017). We explored assemblies built using different approaches: i) the hybrid assembler MaSuRCa v 3.4.2 (Zimin *et al*., 2017); ii) the long reads based assembler Canu 1.8 (Koren *et al*., 2017); iii) WTDBG2 v2.1 (Ruan and Li, 2020) which is also based on long reads; iv) a combination of illumina assembly with platanus v1.2.4 (Kajitani *et al*., 2014) and the DBG2OLC v20180222 (Ye et al., 2016) which uses nanopore data to scaffold the low quality short reads assembly. Each assembly was individually corrected with three iterations of pilon v1.22 (Walker *et al*., 2014). The best assembly was provided by MaSuRCa based on genome size and number of contigs. To obtain a more contiguous assembly Ragout 2.0 was used using the reference strain of *A. fumigatus* found in NCBI (ASM265v1) as template. This improved both assemblies to chromosome level.

### RNA extraction for reverse transcription and real-time quantitative PCR (qPCR)

The mycelia were harvested and washed with cold water followed by freezing in liquid nitrogen and added 1ml TRIzol (Ambion, cat. no. 135405) then stored at −80 L until use. RNA extraction was processed through six cycles of beating in 1 ml ice-cold TRIzol with ∼100 μl volume of silica beads using Bullet Blender (Next Advance). RNA was extracted as previously described (Rio, 2010) and the quality was checked on 2% agarose gel. One µg of total RNAs was subjected to reverse transcription into cDNAs using PrimeScriptTM RT reagent Kit with gDNA Eraser (Takara, cat. no. RR047A) according to manufacturer’s protocol. The resultant cDNA samples were diluted twenty folds and 2 µl of diluted samples were subjected to real-time qPCR analysis as above mentioned.

### RNA purification and preparation for RNA-Seq

Total RNA was extracted as above mentioned. Ten µg of total RNAs was subjected to RNA purification using DNase I as described (NEB, cat. no. M0303) and quality was checked on 2% agarose gel. All RNAs used had a minimum RNA Integrity Number (RIN) value of 7.0. One µg of purified RNAs were used for library preparation. Sequencing libraries were prepared using Illumina NEBNext® Ultra^TM^ Directional RNA Library Prep Kit (NEB, cat. no. 7420) according to manufacturer’s protocol.

### TEs characterization using RepeatMasker

TEs were characterized using RepeatMasker-4.1.1. The script was run for Af293 and CEA17 accordingly (http://www.repeatmasker.org).

### Data mapping and bioinformatics analysis

Raw sequencing reads of histone ChIPseq experiments were quality-checked using FastQC (http://www.bioinformatics.babraham.ac.uk/projects/fastqc/) and aligned to Af293 reference genome (genome version s03-m05-r06) using Bowtie2 (version: 2.2.9) (Langmead, Trapnell, Pop, & Salzberg, 2009). Due to the engagement of H3K9me3 in many repeat regions and the mapped reference matters, we considered all reads of ChIP-seq signal and mapped to each genome reference Af293 (genome version s03-m05-r06) and Af1163 respectively with no mismatches. For H3K9me3 peaks calling, MACS2 was applied using the parameter [macs2 callpeak --nomodel -t ${file} -f BAM -g 29420142 -n ${file}_macs2 --broad --broad-cutoff 0.000001]. If the H3K9me3 peaks cllaed by MACS2 were adjacent, those peaks were manually combined as one peak. The flanking gene lists of H3K9me3 boundary regions were called using in-house scripts <u>(/Users/wangfang/Documents/find_closest_TE_vs_peak.pl)</u>. Gene Ontology (GO) enrichment analysis (Ashburner *et al*., 2000) was performed in FungiDB and were plotted using the online Bioinformatics analysis platform FungiExpressZ (https://cparsania.shinyapps.io/FungiExpresZ/). FPKM for H3K4me3 and RANP II engagement level in gene coding region were called using in-house scripts (<u>/Users/wangfang/Documents/Scripts/learn_plot/zqWinSGR-v4.pl</u>)

### Published RNAseq data mapping

For RNAseq data, raw reads were aligned to *A. fumigatus* af293 reference genome (genome version: s03-m05-r06) using hisat2 (version: 2.1.0) (Pertea, Kim, Pertea, Leek, & Salzberg, 2016) expression level (*e.g.* FPKM) for each annotated gene was calculated using StringTie (version: 1.3.3b) (Pertea *et al*., 2016).

### Mismatch ratio calculation of LINE TE families

The raw sequencing reads of Af293 and CEA17 genomic DNA were aligned to the conserved LINE transposon sequence 592bp, transposon Aft1 and 18S rRNA sequence were used as control (Supplementary Data 5), to calculate the mapping percentage and mismatch reads ratio. If the mapped read contains one or more than one mismatched nucleotide, the read is recorded as mismatched read. The mismatch reads ratio was calculated as the number of mismatched TE/rRNA reads over the TE/rRNA mapped reads to Af293 genome reference.

### High-resolution mass spectrometry analysis

1×10^^4^ spores of each strain were inoculated in 70ml of liquid ANM as previously described and incubated for 72 hours at 37 °C under shaking conditions. The supernatants were frozen dried and 100 mg were extracted with methanol in ultrasonic bath for 40 min. The extracts were then filtered at 0.22u PTFE and submitted to analysis. High-resolution mass spectrometry analyses were performed in a Thermo Scientific QExactive© Hybrid Quadrupole-Orbitrap Mass Spectrometer. Analyses were performed in positive mode with *m/z* range of 100-1500; capillary voltage at 3.5 kV; source temperature at 300 °C; S-lens 50 V. The stationary phase was a Thermo Scientific Accucore C18 2.6 µm (2.1 mm x 100 mm) column. The mobile phase was 0.1% formic acid (A) and acetonitrile (B). Eluent profile (A/B %): 95/5 up to 2/98 within 10 min, maintaining 2/98 for 5 min and down to 95/5 within 1.2 min held for 3.8 min. Total run time was 20 min for each run and flow rate 0.2 mL.min^-1^. Injection volume was 5 µL. MS/MS was performed using normalized collision energy (NCE) of 20, 30 and 40 eV; maximum 5 precursors per cycle were selected. MS and MS/MS data were processed with Xcalibur software (version 3.0.63) developed by Thermo Fisher Scientific.

### Infection of *G. mellonella*

Larvaes of *G. mellonella* in the final (sixth) instar larval stage of development, weighing 275– 330 mg were selected for later experiment. Fresh 2-day conidia from each strain were harvested from ANM plates in PBS solution and filtered through a Miracloth (Calbiochem). For each strain, the spores were counted using a hemocytometer and the stock suspension with 2×10^^8^ conidia/ml was prepeared. The viability of the administered inoculum was determined by plating a serial dilution of the conidia on MM medium at 37 °C. 1×10^6^ conidia (*i.e.* 5 μl) was injected to each larva and 5 μl of PBS was injected at same time as control to observe the killing effect due to physical trauma. The injection was performed by using Hamilton syringe (7000.5KH) via the last left proleg. After infection, the larvae were maintained in petri dishes at 37 C in the dark and were scored daily. Larvae were considered dead by presenting the absence of movement in response to touch.

