## Supplementary figures and images for "Chromatin profiling reveals genome stability heterogeneity in clinical isolates of the human pathogen *Aspergillus fumigatus*"

### Supplementary Figure 1

#### Supplement Figure 1

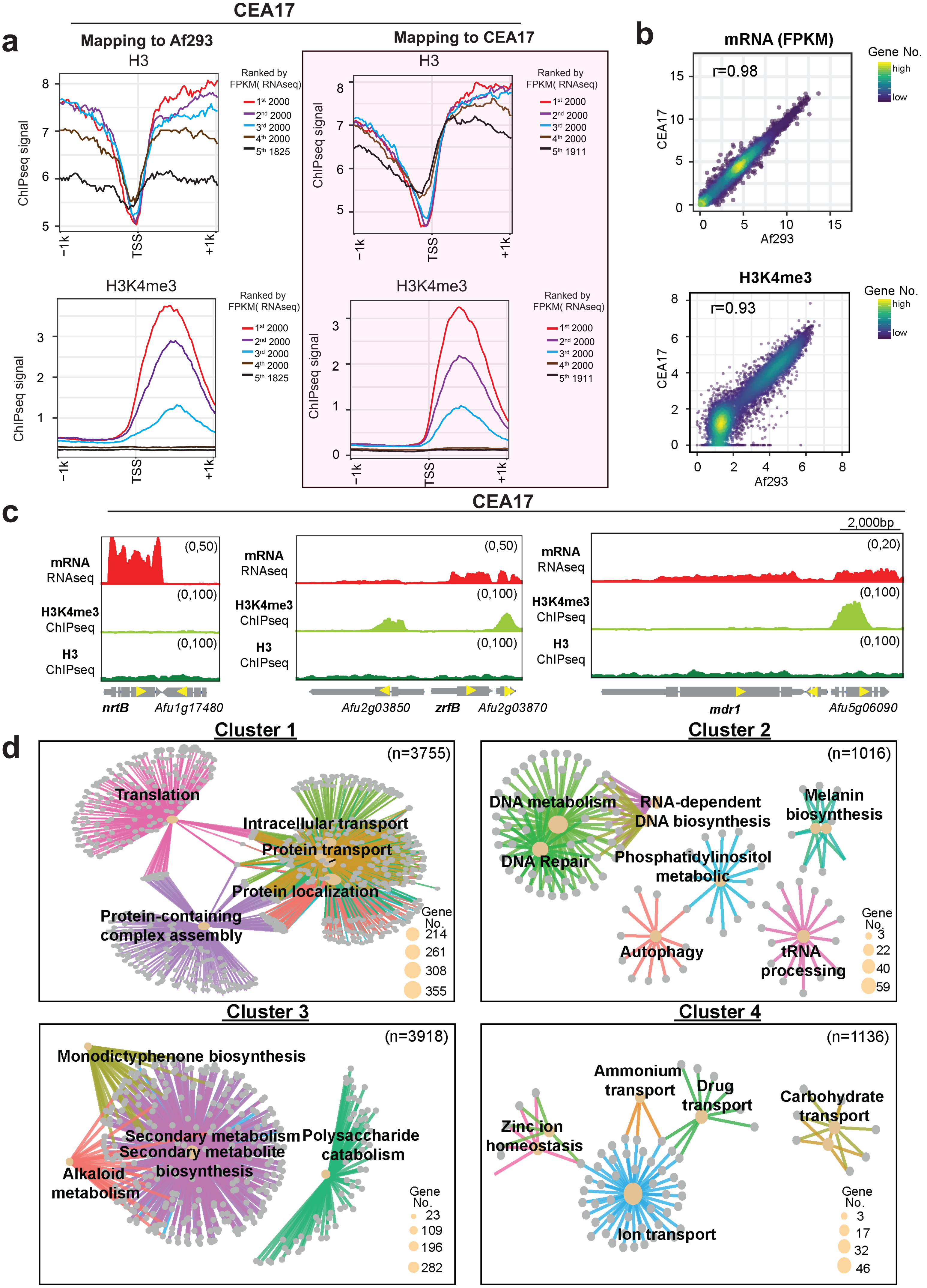

### Supplementary Figure 2

#### Supplement Figure 2

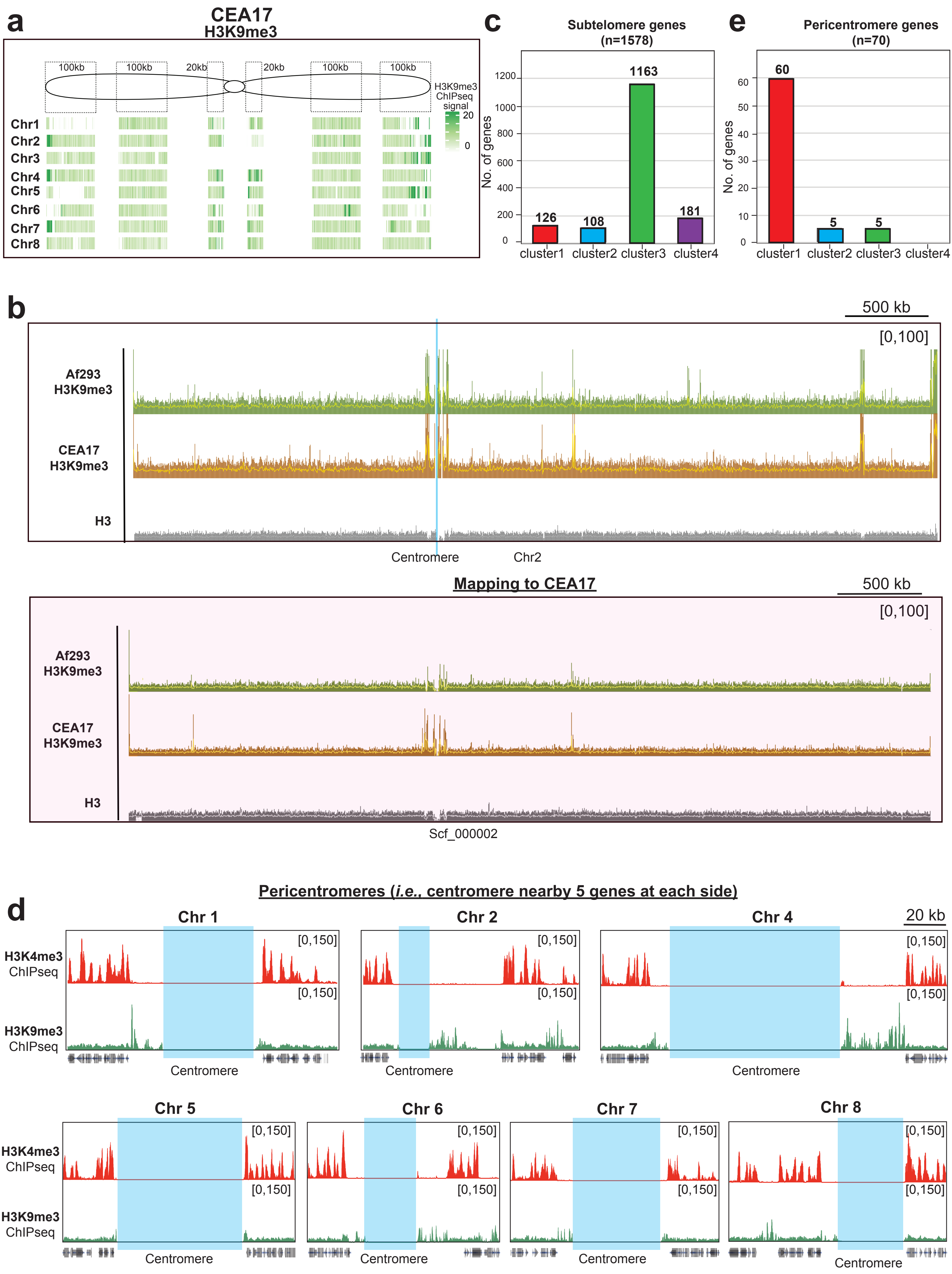

### Supplementary Figure 3

#### Supplement Figure 3

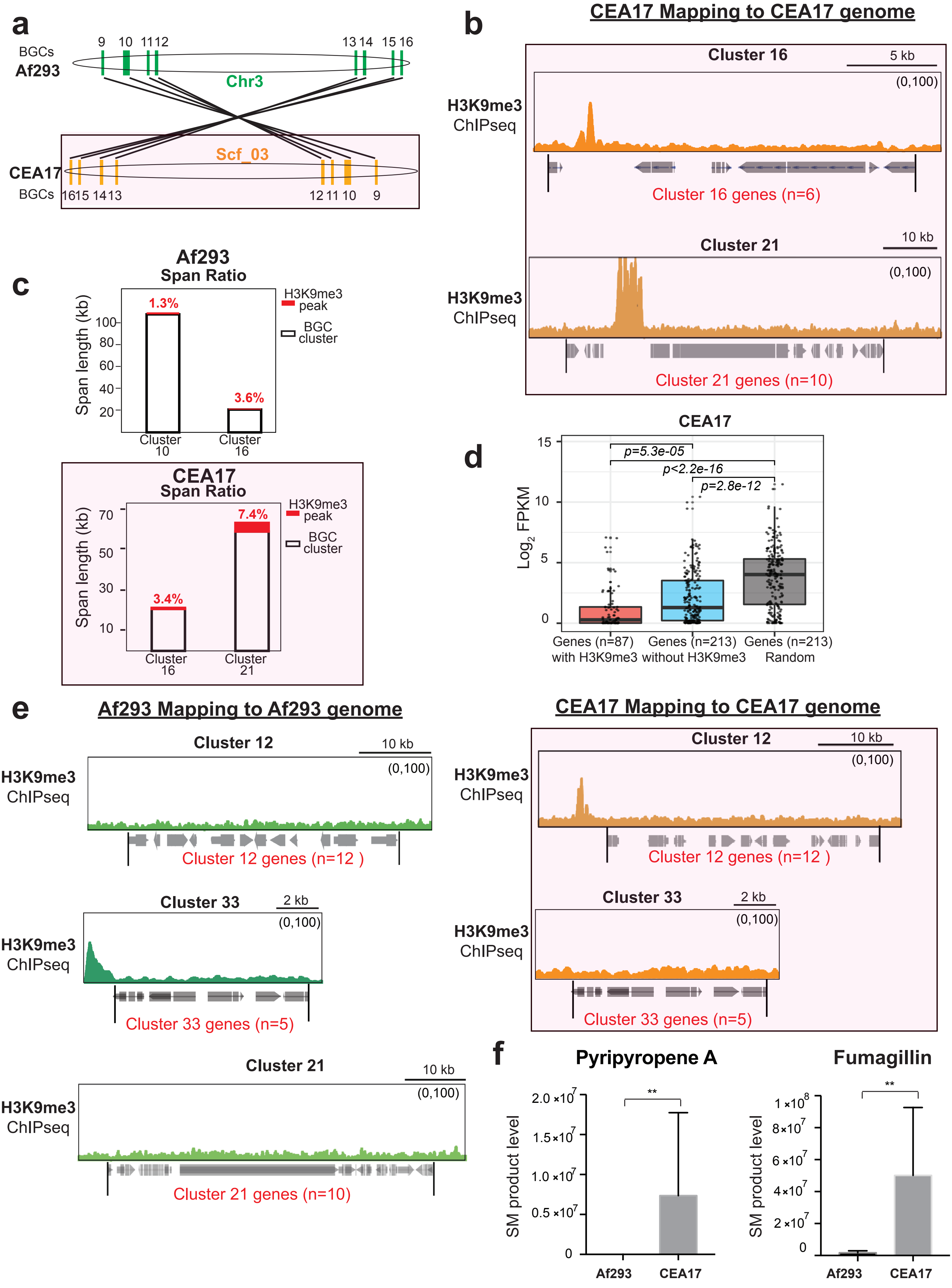

### Supplementary Figure 4

#### Supplement Figure 4

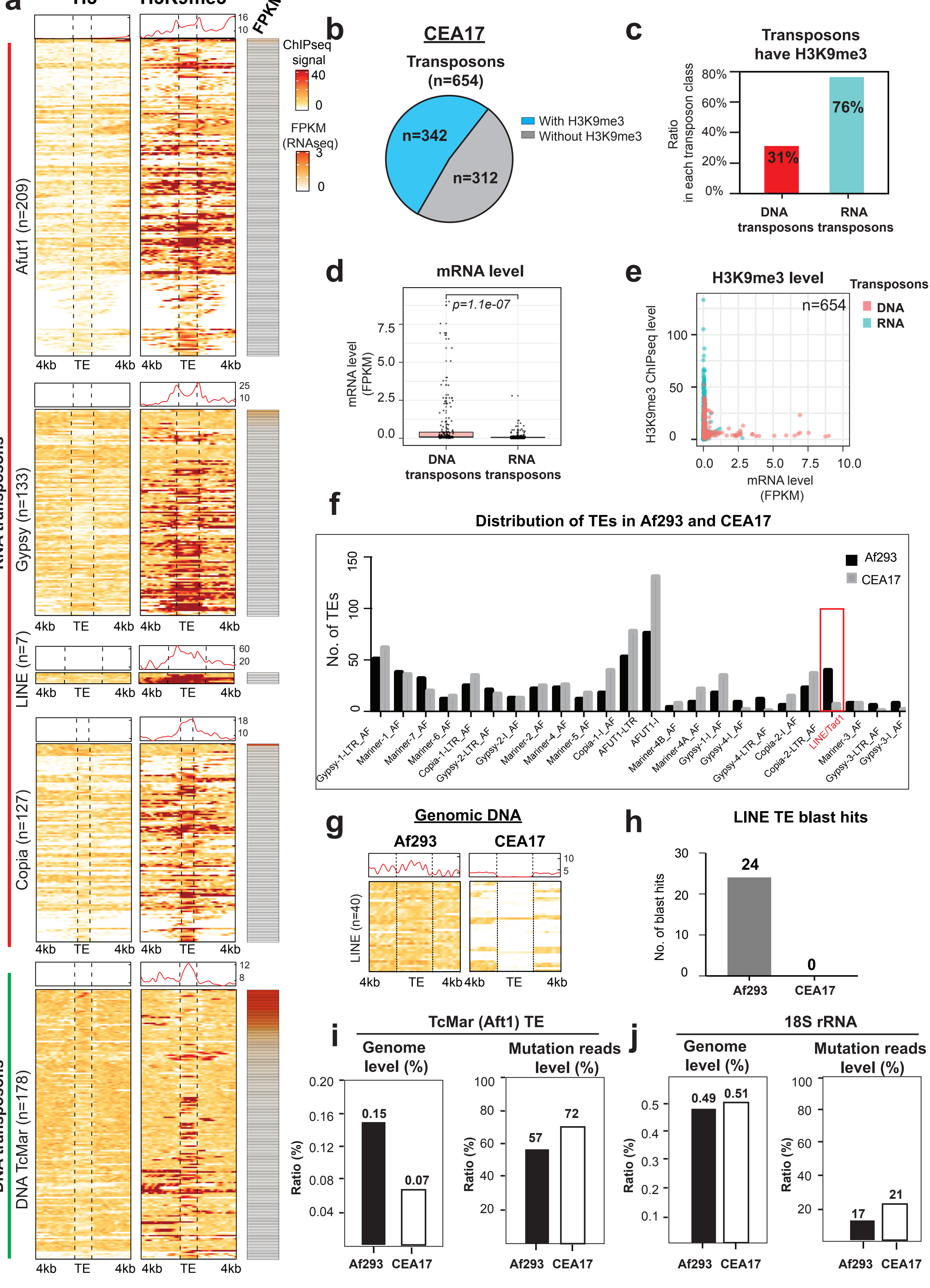

### Supplementary Figure 5

#### Supplement Figure 5

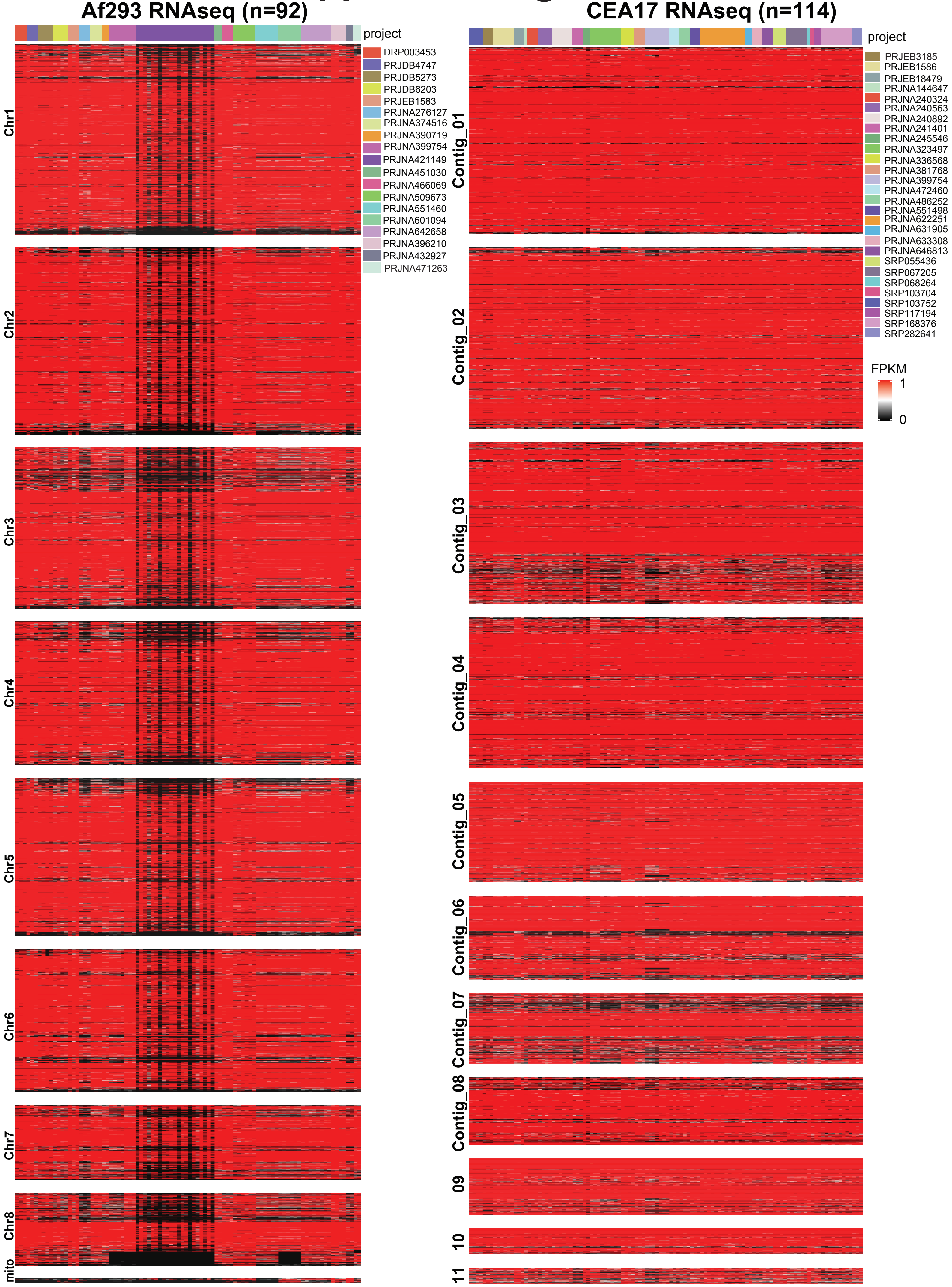

### Supplementary Figure 6

#### Supplement Figure 6

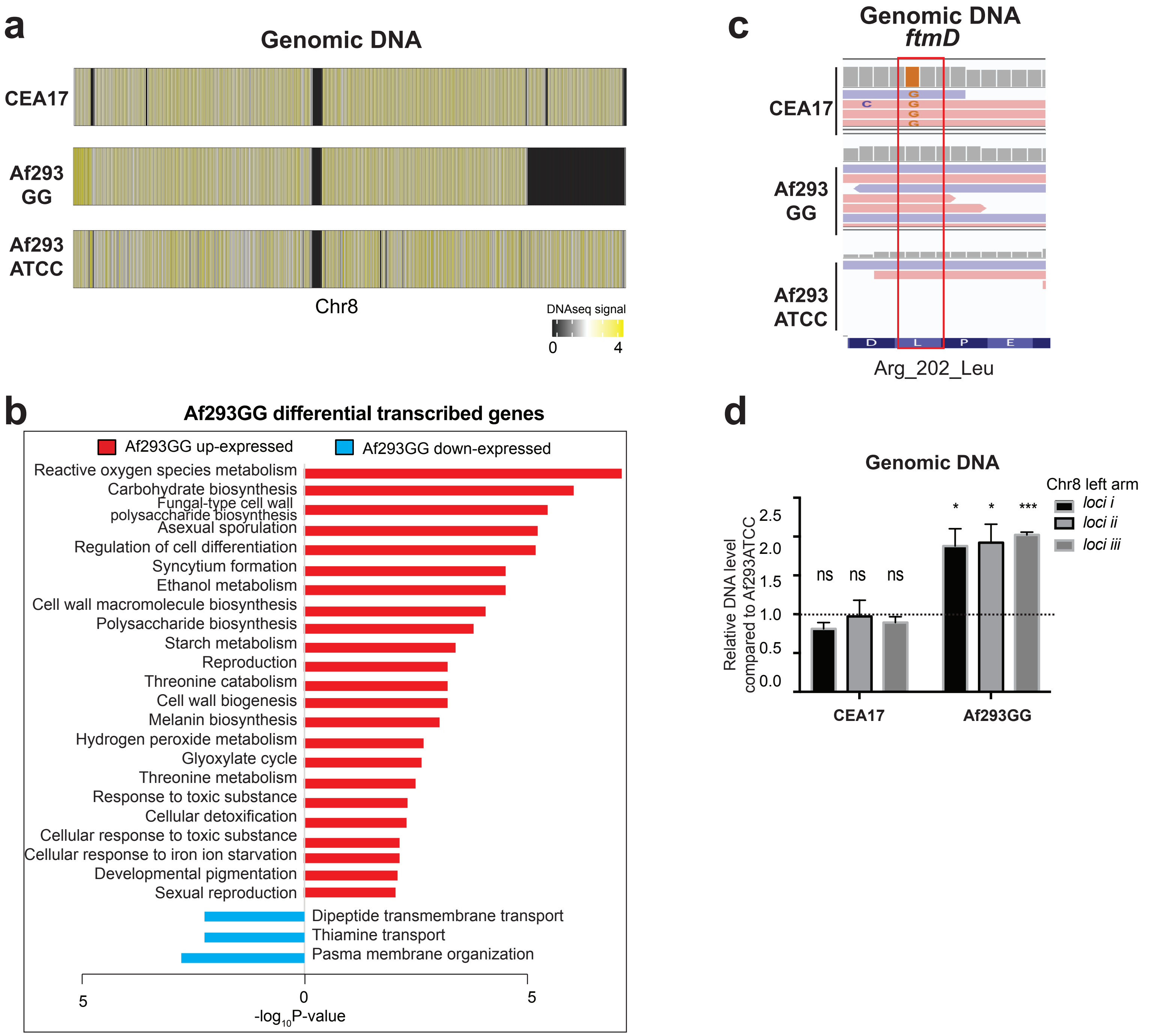

### Supplementary Figure 7

#### Supplement Figure 7

a

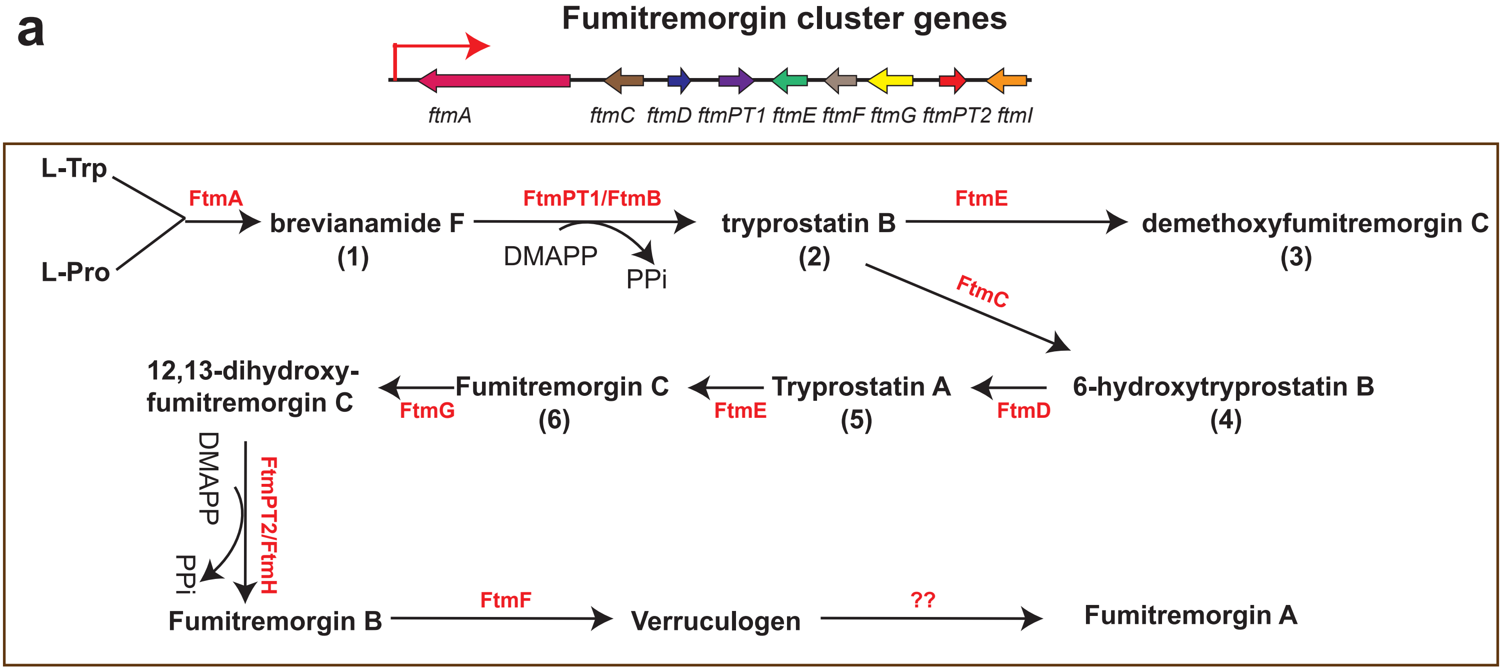


b

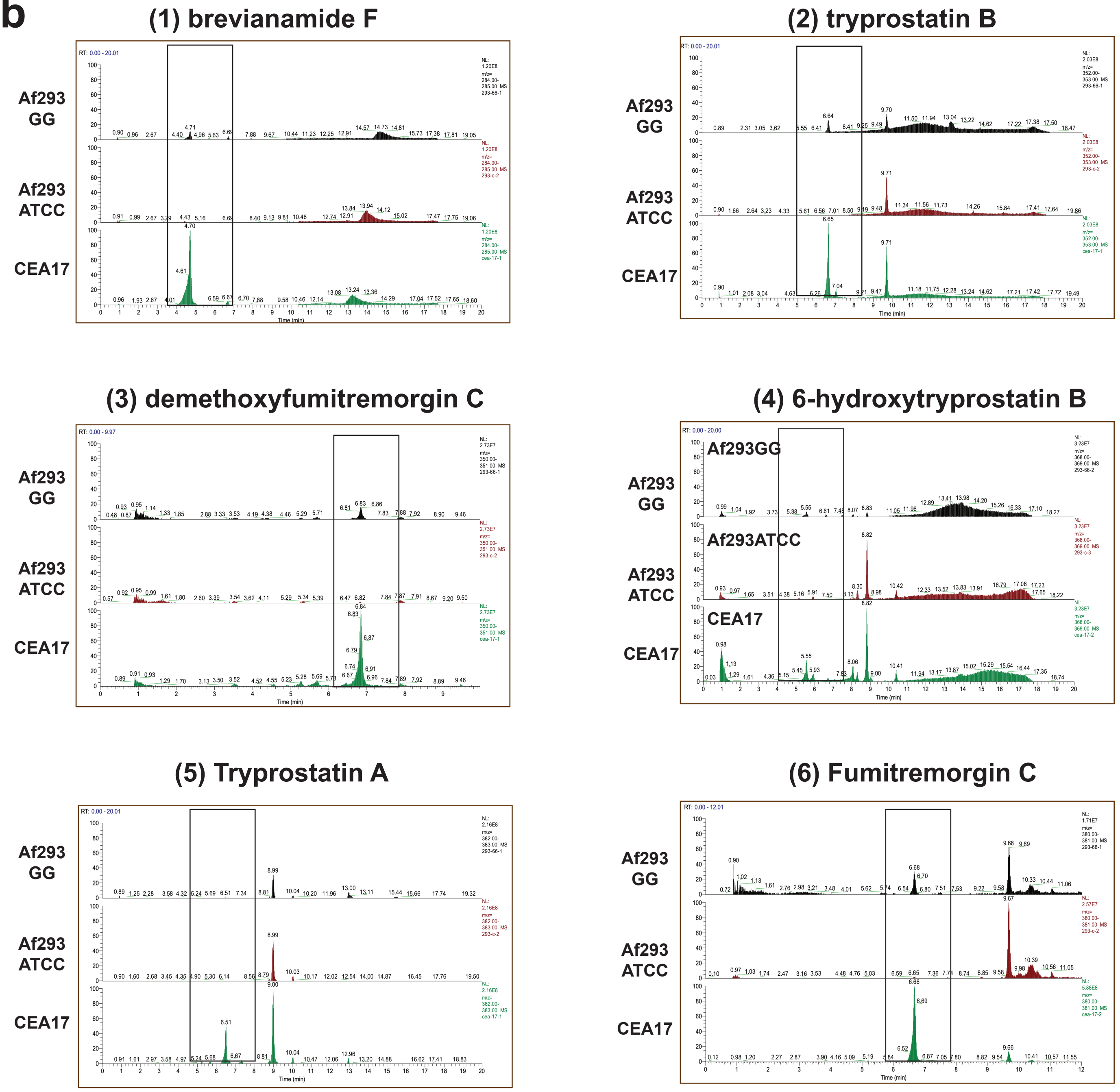

### Supplementary Figure 8

#### Supplement Figure 8

**a**

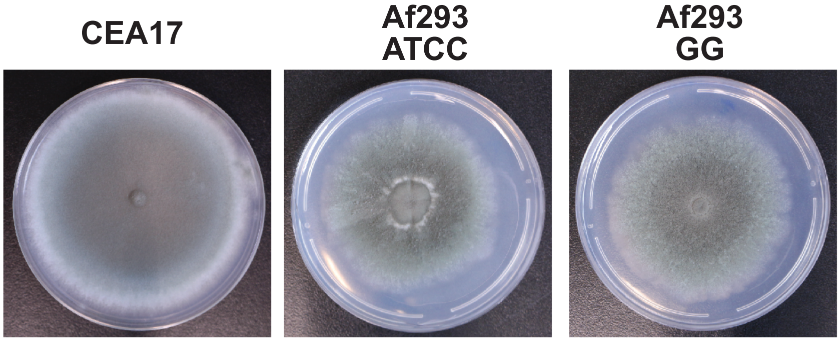


**b**

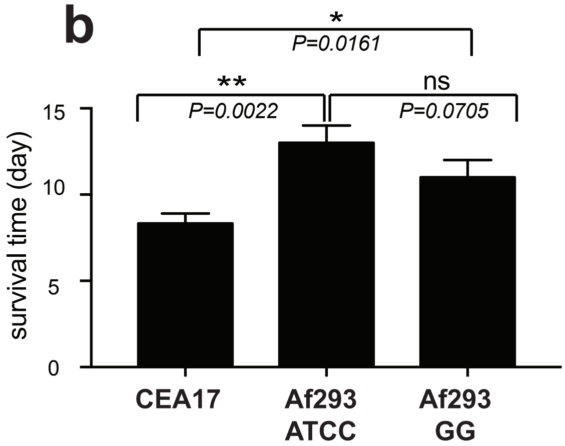
