## Supplementary Text 1 for "Chromatin profiling reveals genome stability heterogeneity in clinical isolates of the human pathogen *Aspergillus fumigatus*"

### **Oxford-nanopore sequencing analysis for two *A. fumigatus* strains**

Table 1: Genome statistics

| Strain | Assembly | Genome size (Mb) | Number of contigs | N50 | Number of Ns |
| --- | --- | --- | --- | --- | --- |
| AF293ATCC | MaSuRCa | 27.6Mb | 76 | 672770 | 100 |
| AF293ATCC | MaSuRCa + Ragout | 29.3 Mb | 8 | 4042076 | 1702405 |
| AF293GG | MaSuRCa | 27.7Mb | 69 | 945918 | 0 |
| AF293GG | MaSuRCa + Ragout | 28.6 Mb | 8 | 3906814 | 1226787 |
| Reference genome | - | 29.4 Mb | 8 | 3948441 | 575000 |

#### **Detection of variations in chromosome VIII:**

We compared both assemblies to the reference genome and observed the deletion of the right hand part of chromosome VIII in Af293GG (see figure 1). Images were generated with lastal (Kiełbasa *et al*., 2011) (v941) and circos v0.69-8 ( Krzywinski *et al*., 2009). To ensure it was not an effect of the fragmentation of the genome, we mapped the illumina reads of both strains to the reference genome using BWA v 0.7.17-r1188 (Li and Durbin, 2009) and samtools v1.9 (Li *et al.*, 2009) and observed the same deletion using IGV v2.8.2 (Robinson *et al*., 2011) (see figure 2). The duplication of the beginning of the left arm of chromosome VIII was also observed as an increase of coverage in the read mapping.

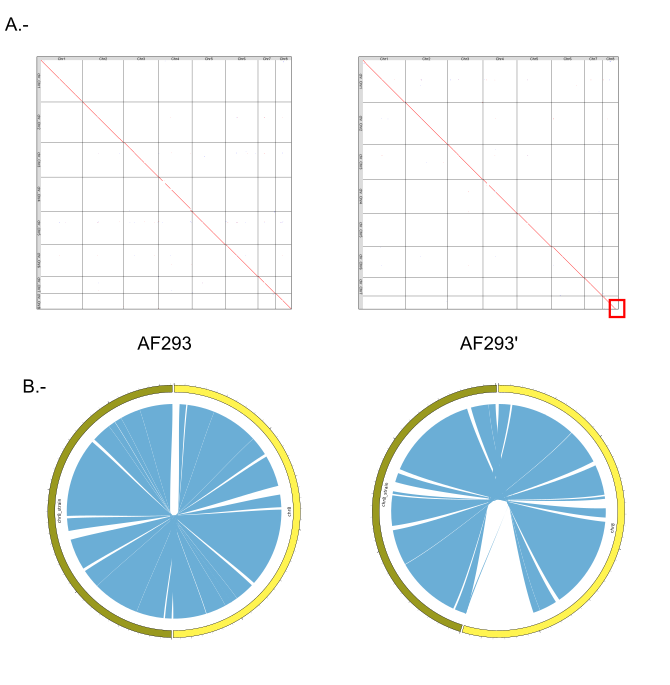

Figure 1.- A.- Dot-plot image of the genomes of AF293ATCC (AF293) and AF293GG (AF293’) against the *A. fumigatus* reference genome. On the left is the plot for AF293ATCC and on the right the plot for AF293ATCC missing a piece of chromosome VIII. B.- Circos plot of the comparison chromosome VIII of AF293ATCC (left) and AF293GG (right) against the reference.

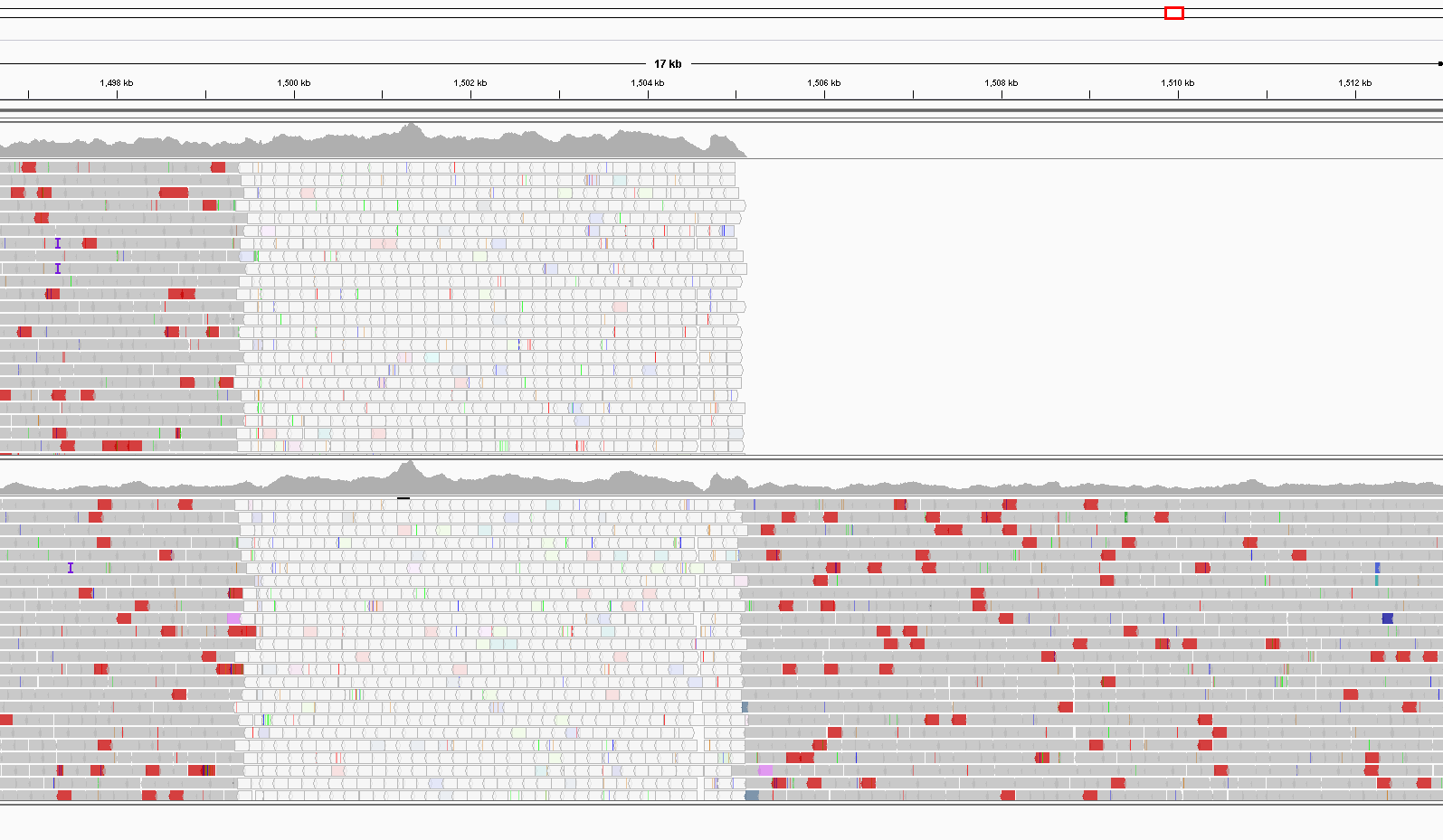

Figure 2: Snapshot of IGV viewer at the point where the AF293GG genome (upper track) has the deletion. For comparison, the lower track contains the read mapping for AF293ATCC.

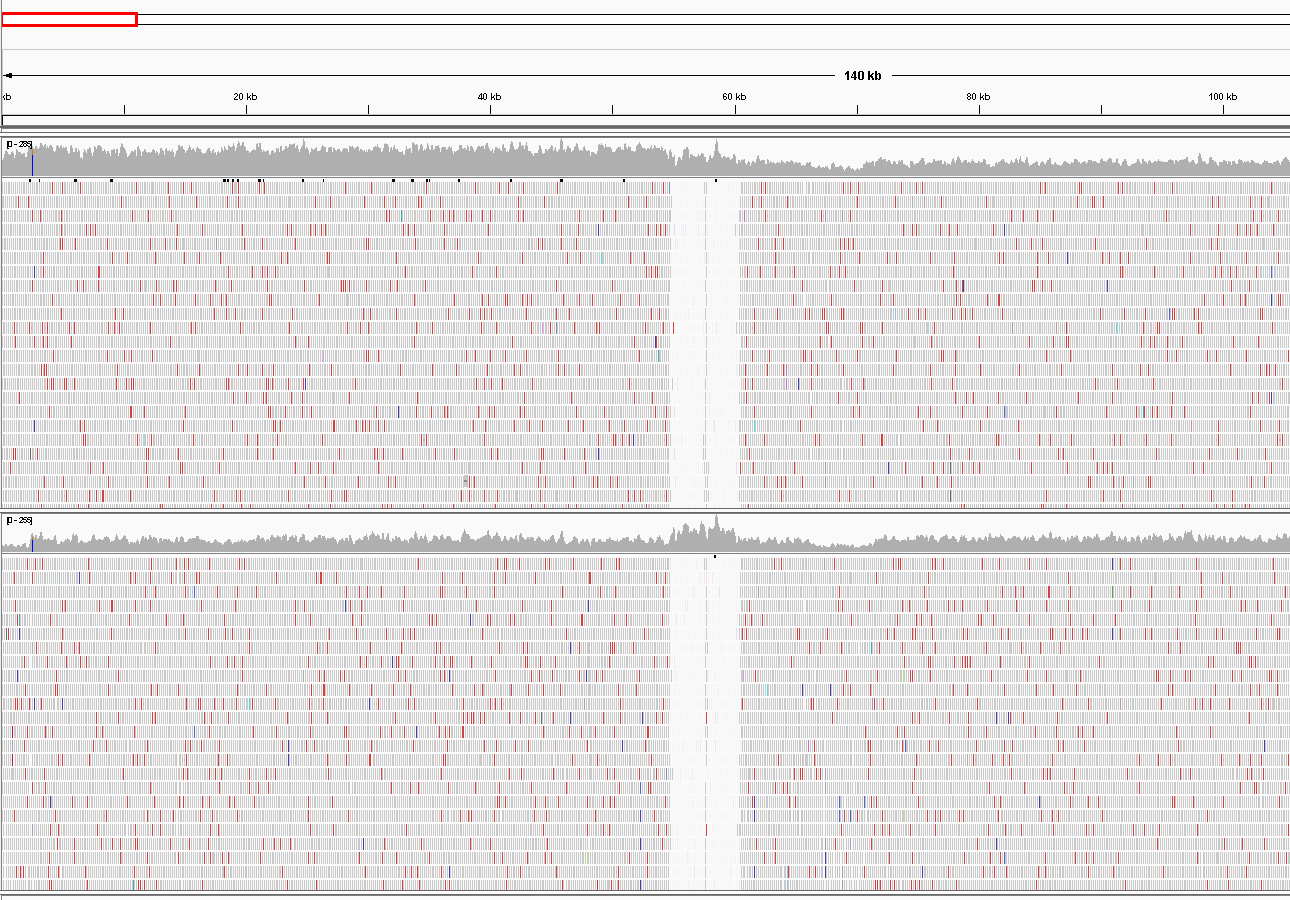

Figure3: Snapshot of IGV viewer of the duplication in the left arm of chromosome VIII in AF293GG (upper track). The lower track contains read mappings for AF293ATCC for comparison.

### **Results**

We assembled the genomes of AF293ATCC and AF293GG to chromosome level assemblies using a hybrid assembly strategy and with a final scaffolding step relying on the *A. fumigatus* AF293 strain stored in NCBI (ASM265v1) (see methods). We used these assemblies to confirm the re-arrangements noted in the manuscript. We found a deletion at the end of chromosome VIII in strain AF293GG. To ensure that this was not related to the low coverage present in the illumina and Nanopore data we also mapped reads directly to the reference genome, which verified the presence of a deletion at the right hand side of chromosome VIII starting at position 1,505,138 after what appears to be a repetitive region. Indeed, in positions 1499830 to 1503369 a transposable element from the LINE family was identified in the reference (see Figure 2). A total of 106 coding genes were lost with this deletion. Additionally a duplication was observed at the left arm of chromosome VIII, spanning a region of roughly 60 kb, where 20 coding genes were located. This region was detected due to the presence of a doubled region in terms of read coverage but was otherwise absent in the assembled genome indicating it had been collapsed (see Figure 3).
